# *In vivo* targeted and deterministic single cell malignant transformation

**DOI:** 10.1101/2024.01.30.577941

**Authors:** Pierluigi Scerbo, Benjamin Tisserand, Marine Delagrange, Héloïse Debare, David Bensimon, Bertrand Ducos

**Affiliations:** Laboratoire de Physique de l’Ecole Normale Supérieure LPENS, ENS, PSL Research University, CNRS, Sorbonne Université, Université de Paris, 24 rue Lhomond 75005 Paris, France; High Throughput qPCR Core Facility of the ENS, Ecole Normale Supérieure, PSL Research University, IBENS, 46 rue d’Ulm 75005 PARIS; Dept. Chemistry and Biochemistry, UCLA, 607 Young Dr.E, Los Angeles 90095; Inovarion, 75005 Paris, France

## Abstract

Why does a normal cell possibly harboring genetic mutations in oncogene or tumor suppressor genes becomes malignant and develops a tumor is a subject of intense debate. Various theories have been proposed but their experimental test has been hampered by the unpredictable and improbable malignant transformation of single cells. Here using an optogenetic approach we permanently turn on an oncogene (KRASG12V) in a single cell of a zebrafish brain that, only in synergy with the transient co-activation of a reprogramming factor (VENTX/NANOG/OCT4), undergoes a deterministic malignant transition and robustly and reproducibly develops within 6 days into a full-blown tumor. The controlled way in which a single cell can thus be manipulated to give rise to cancer lends support to the “ground state theory of cancer initiation” through “short-range dispersal” of the first malignant cells preceding tumor growth.

## INTRODUCTION

How cancer arises from a single normal cell is still the subject of active debate, affecting intervention strategies. While many cells may harbor oncogenic mutations, only a few unpredictably end-up developing a full-blown tumor^1,2,3,4^. Various theories have been proposed to explain that transition^5,6,7,8^, but none has been tested *in vivo* at the single cell level. Cancer initiation is thus believed to be a *rare* event taking place at the level of *individual cells*^9,10,11,12,13^ arising as a result of the accumulation of genetic mutations in so called Mut-driver genes (oncogenes such as KRAS^14,15^, tumor suppressor genes^16^ such as TP53 or life-span^17^ genes such as TERT). Notably, mutant KRAS is the most frequent driver of several cancers^18,19^: about 27% of all human cancers, 45% of colorectal and 90% of pancreatic cancers^20^.

Recently, it has been shown that genes involved in embryonic development, pluri/multipotency and cell reprogramming such as VENTX/NANOG and POU5/OCT4 are abnormally reactivated in late cancer stages, where acting as Epigenetic Drivers (Epi-Drivers) they empower cancer cells with Cancer Stem Cell (CSCs) features, resistance to anti-cancer therapies and potential for cancer recurrence/relapse^21,22,23,24,25,26,27^. Although it is evident that such Epi-Drivers confer a selective advantage to CSCs in a full-blown cancer, whether they play a role during the early phases of malignant transformation is still unknown.

Current approaches to the study of cancer use constitutive or conditional expression of Mut-driver (or Epi-driver genes^28^) in specific tissues, i.e. in many cells, even though only a small subset of these cells eventually leads to the growth of tumors^29^, often observed when the tumor already consists of many thousands of heterogeneous abnormal cells. Hence, carcinogenetic processes observed among sibling organisms^30,31,32^ occur with variable latency period from the onset of induction, at different locations and develop asynchronously. As a result, the initial stages of tumorigenesis are difficult to study, the state of the cell(s) of origin difficult to assess and control while its cellular environment is perturbed by the induced expression of the Mut-or Epi-driver genes. Due to the rarity of the event *in vivo*^32^, a statistically relevant single-cell tracking and characterization of the early stages of tumorigenesis has therefore never been done. This emphasises the need to predict or control the cell undergoing malignant transformation *in vivo* in order to pave the way for a study of the cellular and molecular events involved in the initial stages of tumorigenesis.

To address these issues, we have developed an optogenetic approach to control the expression of an oncogene in a single cell^33^, with the goals of: (1) measuring the probability of the malignant transformation of a single cell in a live organism in various backgrounds and (2) tracking and characterising the development of a tumor from the original cell. **In the following we show that the synergy between only two factors: the oncogene kRasG12V and a reprogramming factor (Ventx, Nanog or Oct4) increases the probability of carcinogenesis from a single cell by many orders of magnitude when compared to the expression of either of these genes (or none).**

Optogenetics approaches allow for the photocontrol and monitoring of the activity of biomolecules *in vivo*. The approach we developed uses a photoactivable analog of tamoxifen (caged cyclofen, cCYC) to control the activity of proteins fused to the ERT-receptor^34,35^ (a modified estrogen binding domain^36^) (Fig. 1A). These protein constructs are sequestered by cytoplasmic chaperones. Once cCYC is uncaged by light (with one-photon illumination at ∼375 nm or two-photon at ∼750 nm), cyclofen (CYC) is released^34^. It binds to the ERT-receptor and releases the fused protein (e.g. a Cre-ERT recombinase) from its complex with cytoplasmic chaperones^35^ (Fig. 1A).

**FIGURE 1:**
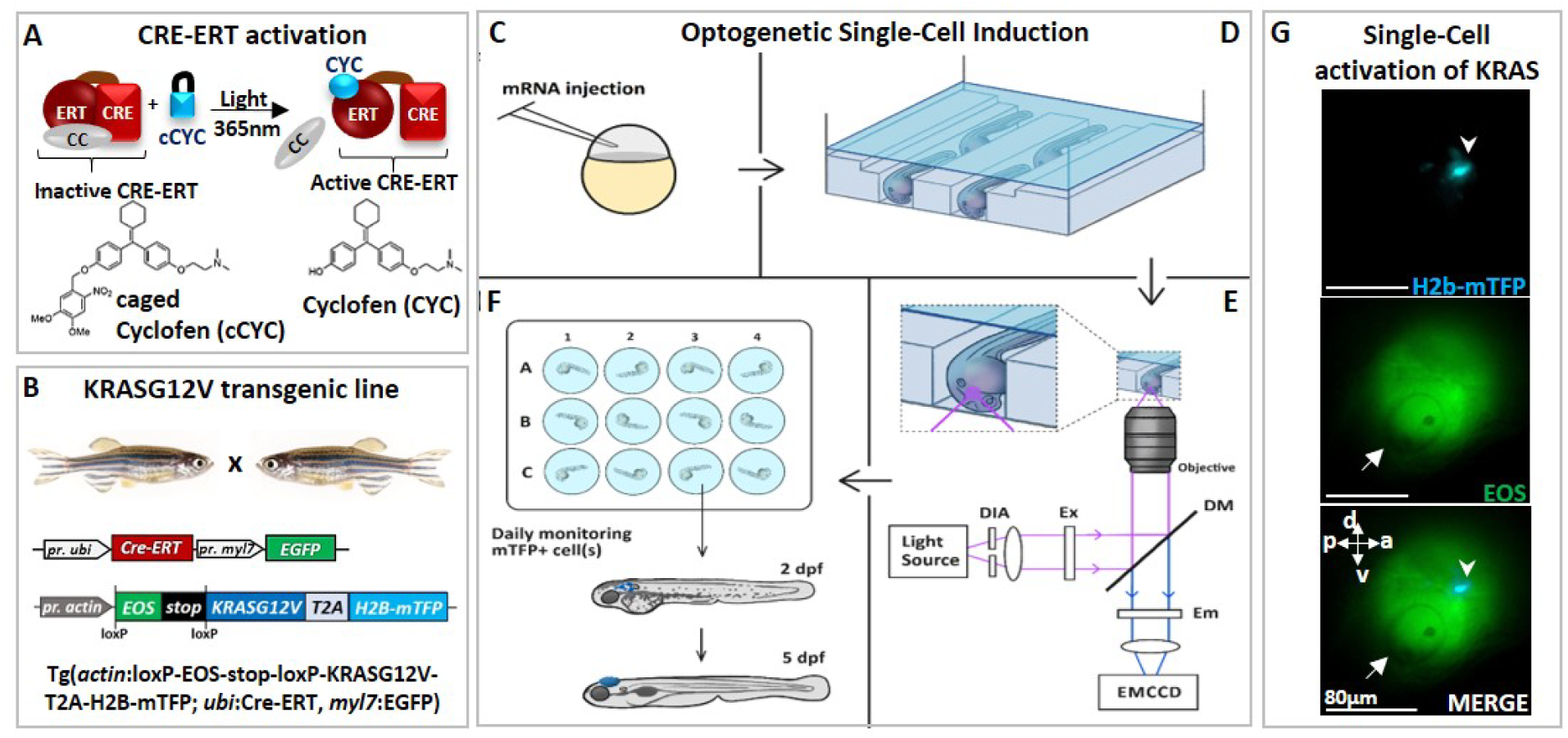
Optogenetic set-up for single cell induction. **(A)** Photo-control of a protein fused to an Estrogen Receptor (ERT) is achieved by releasing the protein from its complex with cytoplasmic chaperones (CC), upon uncaging of cCYC. **(B)** The transgenic zebrafish line engineered to express an oncogene (KRASG12V) upon photo-activation of a CRE-recombinase fused to ERT (CRE-ERT) (as shown in A). **(C)** The mRNAs of Ventx-GR and mRFP (used as a marker) are injected at the one cell stage. **(D)** At 1dpf the embryos are mounted in channels in an agarose gel and incubated for 45 min in cCyc on a microscope stage. **(E)** They are washed and illuminated at 405nm on a microscope to uncage cCyc close to the otic vesicle. A diaphragm (DIA) defines an illumination zone of ∼80μm diameter (see G). Excitation (EX), Dichroic mirrors (DM) and emission (EM) filters allow for visualization of Eos and mTFP. The cell in which KRASG12V has been induced is observed within ∼1h by the fluorescence of mTFP (see G). **(F)** The embryos are transferred into individual wells, incubated overnight in DEX, washed at 1dpi and monitored over the next 5 days. **(G)** Lateral view of zebrafish at ∼1h post-activation displays a single induced cell (top: blue spot shown by arrowhead in mTFP channel) in the illumination region (middle: Eos channel) in the vicinity of the otic vesicle (white arrow) and bottom: merger of both channels. Body axes (a: anterior; p: posterior; d; dorsal; v: ventral) are shown.

## RESULTS

We used this approach to photocontrol the activity of a Cre/loxP recombination system in a transgenic zebrafish line (Tg(*actin*:loxP-EOS-stop-loxP-KRASG12V-T2A-H2B-mTFP; *ubi*:Cre-ERT; *myl7*:EGFP) carrying a floxable (loxP flanked) EOS gene (coding for a green fluorescent protein) upstream from the oncogene^33,34^ (KRASG12V). In such a transgenic zebrafish line hereafter referred as KRASG12V line (Fig.1B, Fig.S1B) the expression of EOS can be switched to KRASG12V and H2B-mTFP (nuclear blue fluorescence) upon 1h incubation in cCYC, followed by washing and UV (365 nm) cCYC uncaging which triggers CRE-ERT activation (Fig. 1C); mTFP fluorescence can be observed about 30min post-illumination (Fig. S1B) and is stably maintained in zebrafish (Fig.1G, Fig. S1C). Consistent with an acquired refractory cell state and a loss of oncogenic competence^30^, whole body expression of the oncogene at 1dpf did not result in tumorigenesis. Notice that consistent with previous studies^33–35^, where cCYC stability has been validated both *in vitro* (by chemical analyses), and *in vivo* (with fluorescent reporters of uncaging), in absence of illumination no spontaneous uncaging of cCYC and thus activation of the oncogene (i.e. blue fluorescent cells) is observed. Notice also that once cCYC is uncaged embryos are maintained in E3 medium with no cyclofen and thus no activation of the oncogene.

To test for the possible impact of a reprogramming factor on the malignant transition, embryos from this transgenic line were injected at the one-cell stage with the mRNA of a construct consisting of a glucocorticoid receptor (GR) fused with a reprogramming factor (Ventx, Nanog or Oct4)^24,37^ (Fig.S2). The resulting protein (e.g. Ventx-GR) is sequestered by cytoplasmic chaperones and transiently activated upon incubation of zebrafish in Dexamethasone (DEX) (Fig. S2A). Activation of the oncogene^33^ at 1dpf, if and only if followed by transient activation of Ventx (or Nanog or Oct4) did yield reproductible hyperplasic outgrowths in many different tissues (including brain, intestine, pancreas and liver; Fig. S3B-E). Immuno-histochemistry detection of phospho-ERK activity (Fig. S4A), Hematoxylin & Eosin staining (Fig. S4B) and RT-qPCR of selected genes (Figs. S5; S6A,B) all display features associated with tumorigenesis. None of the controls (activation of kRasG12V only, Ventx-GR only, incubation in cyclofen or dexamethasone, etc.) developed hyperplasia, but rather grew into normal zebrafish (Figs. S1B,C; S3A). These controls imply that the observed tumor is only a result of the joint activation of kRasG12V (permanently) and a reprogramming factor (transiently).

Since both kRasG12V and Ventx have been reported to play a role in brain cancer^38^, we decided to activate the oncogene kRasG12V at 1dpf in a single normal cell of the brain of a transgenic zebrafish (injected with the mRNA of Ventx-GR at 1 cell stage). Activation of the oncogene in a single cell was achieved by 1h incubation in cCYC, followed by washing and illumination at 405nm to uncage cCyc in a small area (diameter ∼80 μm) of the brain (in the vicinity of the otic vesicle) (Fig. 1G). Notice that following uncaging the embryos are maintained in a cyclofen free E3 medium.

Subsequent to cCYC uncaging, with probability ∼50% the oncogene is activated in a single cell of the illuminated area (activation statistics is shown in Fig.2A), identified within ∼1h by the blue fluorescence of its nuclear marker (H2B-mTFP), see Figs. 1G, 2A, S7. H2B-mTFP positive cells are only observed at the site of illumination (i.e. brain), attesting to the precision of our optogenetic method (Fig. 2A). This observation specifies whether one, two or more cells were activated (Fig. S7) within ∼1h post-illumination. Notice that activation of the oncogene alone does not give rise to cancer (the activated cell usually disappears within a few days, Fig. S10A), which implies that the proliferation of kRasG12V positive (mTFP^+^, blue fluorescent) cells - reported next - is not due to late activation of the oncogene (and its fluorescent marker).

**FIGURE 2:**
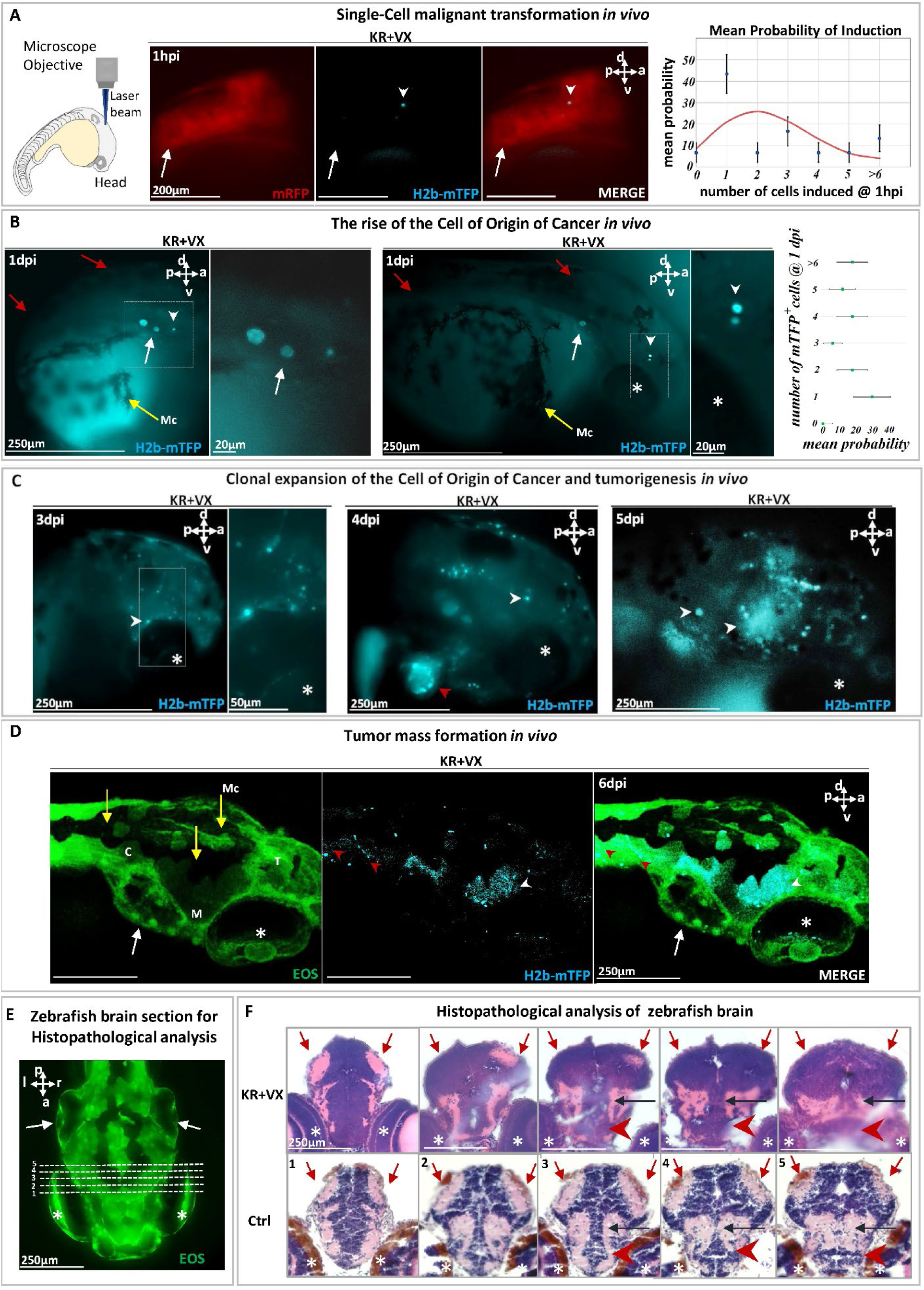
Malignant transformation of a single cell triggering carcinogenesis *in vivo.* **(A)** At 1dpf, a single cell in a zebrafish brain was photo-induced to express the oncogene KRASG12V, identified (white arrowhead) within ∼1h by the blue fluorescent H2B-mTFP. Membrane-bound mRFP is used as tracer. Transient (24h) DEX activation of Ventx is done following photoactivation. The probability *p* of inducing one or more (blue fluorescent) cells is shown on the right panel, together with the Poisson distribution (red curve) expected for the independent induction of *k* cells (error bars are statistical errors on the mean: 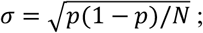 where *N* is the total number of observed embryos). **(B)** At 1day post induction (1dpi), the activated cell may have divided (middle panel) giving rise to two mTFP^+^ (blue fluorescent) cells (white arrowhead) or may have not divided (left panel). The probability of observing *k* blue fluorescent cells at 1dpi is shown on the right panel. **(C)** At 3dpi the original cell expanded clonally (white arrowheads) by short-range dispersal within the brain. At 4dpi the brain has been colonized (middle panel) by the progeny of the activated cell that display tumor growth as well as dispersal in the head or entering into the cardiovascular system (red arrowhead). At 5dpi (right panel), a tumor mass is formed. **(D)** Confocal microscopy of a larval head displaying tumors (white arrowheads) and dispersal in the trunk (red arrowheads). **(E)** Histopathological sections of larval brain (dorsal view). Dashed lines (1 to 5) indicates sections showed in **(F)** Hematoxylin & Eosin (H&E) staining of brain at 5dpi of KR+VX-induced larva (depigmented) is compared to normal brain (Ctrl, melanocytes in brown). At 5dpi, the optic tectum (red arrows), the tegmentum (black arrow) and hypothalamus (red arrowhead) are infiltrated by a dysplasic tumor, progeny of the initial induced single cell. An asterisk (*) indicate the eye and a white arrow the otic vesicle. Scale bars and the body axes (a: anterior; p: posterior; d; dorsal; v: ventral) are shown. T=telencephalon; M=Mesencephalon; C= Cerebellum; Mc= Melanocytes (yellow arrows).

Indeed, if and only if following the local expression of the oncogene, Ventx-GR is transiently activated (by 24h incubation in dexamethasone) does the cell divide and proliferate (Figs. 2B-C, S8, S9). We observed that at 1-day post induction (dpi) ∼50% of the induced cells had divided and expanded clonally by a factor ∼3 (Fig. 2B). In the other 50% of cases, the activated single cell had neither divided nor died (Fig. 2B). Surprisingly, at 3-5 dpi, we observed in all zebrafish larvae (n=30) that the induced cell(s) gave rise to progeny that display short-range dispersion (Fig. S9A, B) and then give rise to a tumor mass in the brain with local infiltration of malignant cells (Fig. 2C) Hematoxylin-Eosin (H&E) staining of the brain tissue (Fig. 2D, E) and metastases (Fig.3) further confirm the malignant state of the induced cell and its progeny.

**FIGURE 3:**
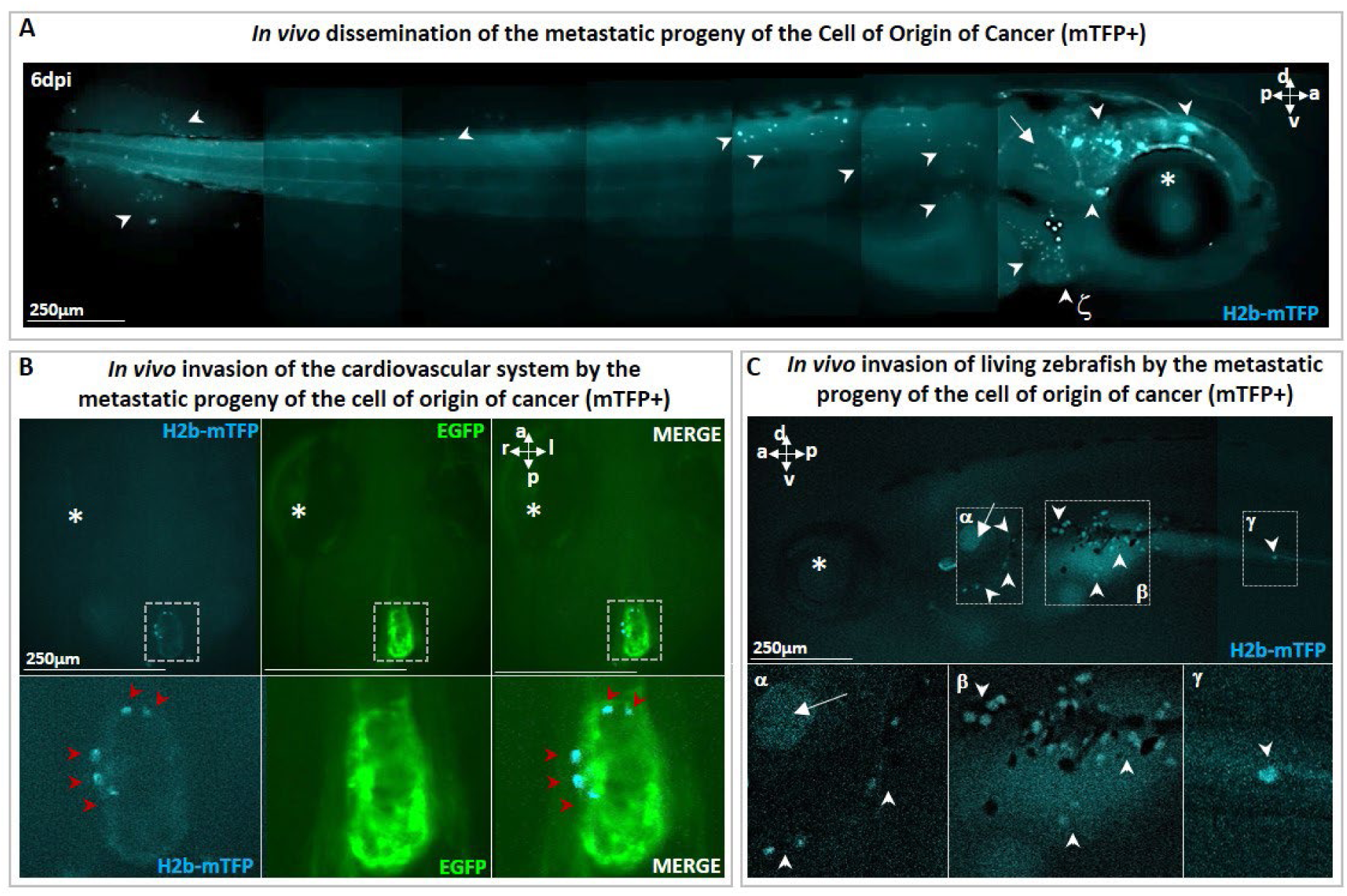
Metastatic cells following single cell malignant transformation. **(A)** A zebrafish larva, in which a single cell in the brain was photo-induced (at 1dpf) to express the oncogene KRASG12V by the blue fluorescence of the expression marker (H2b-mTFP) as in (Fig. 2), show that the progeny of the Cell of Origin of Cancer give rise to a tumor mass in the brain (white arrowhead), as well as to migrating metastatic-like cells that disseminate in the whole organism (white arrowhead), some localizing in proximity of arterial branchial arches (indicated by ζ and white arrowhead), trunk and tail fin. **(B)** H2b-mTFP positive cells can migrate far from the site of induction (brain) and colonize ectopic tissues located in the heart and **(C)** the bottom of otic vesicle (in proximity of the primary head sinus, designated by α), the digestive tract (designated by β and γ) a feature characteristic of metastatic cancer cells. Note that no anaesthetics (e.g. tricaine) or mounting media (low melting point agarose or methylcellulose) were used to block live zebrafish, during both monitoring and imaging performed on live zebrafish. An asterisk (*) indicate the eye and a white arrow the otic vesicle. Scale bars and body axes (a: anterior; p: posterior; d; dorsal; v: ventral) are indicated.

In parallel with tumor mass formation in the brain (Fig. 2), we observed that some cells of the progeny re-localized to new loci far from the brain, such as the heart (Fig. 3A, the digestive tract (Fig. 3B, Fig. S6D) and the trunk (Fig. S6C, Fig.S8B). We therefore deduce that the transient activation of Ventx alters the state of a cell expressing a mutated oncogene (i.e. KRASG12V) and induces its malignant transformation *in vivo*, with tumorigenic potential and the capacity to generate invasive progeny. Notice that metastatic cells must be the progeny of the initial (blue fluorescent) cell as the larvae are incubated in embryo medium with no cCYC. Furthermore, **absent VentX activation we do not observe proliferation** and metastatic behaviour, rather the initially induced cell disappears, see Fig.S10A.

Of the larvae in which a few (1-6) cells were expressing the oncogene and in which Ventx was transiently activated (KR+VX cell), all (N = 30) developed tumors within 5 dpi. The frequency of tumor development is therefore F_1_=1. Conversely the probability for such a cell to not develop a tumor is F_0_=0. Due to the finite size of the sample, we estimate the probability of tumor development from a single cell to be: P_1_ > 80% (χ^2^ = 3,75; df=1).

To definitely confirm the carcinogenic nature of the KR+VX induced cells we isolated and injected a single cell from a hyperplasic tissue (identified by the blue fluorescence of its nucleus) into a naïve host zebrafish larva. This led to integration, migration and colonization of the host tissues by the progeny of the transplanted cell (Fig. 4A,B), with strong pERK activity detected in tumor masses of the host (Fig. 4B). Out of 52 transplanted host zebrafish, 31 developed tumors, a probability of tumor development (60%) consistent with previously reported efficiency of tumor cells transplantation in zebrafish^39^. The capacity of the progeny of the induced cells to metastasize and develop a tumor mass when transplanted in a naïve zebrafish is - to the best of our knowledge - the **operational definition of a malignant tumor**. This result definitely confirms the cancerous nature of cells expressing a mutated oncogene and exposed to the transient activation of Ventx, with subsequent alteration of the homeostasis. Notice that injection into a naïve larva of an oncogene expressing cell (identified by its nuclear blue fluorescence) which **did not** experience the transient activation of a reprogramming factor **does not** yield a tumor and disappears within a few days (Fig. S10B).

**FIGURE 4:**
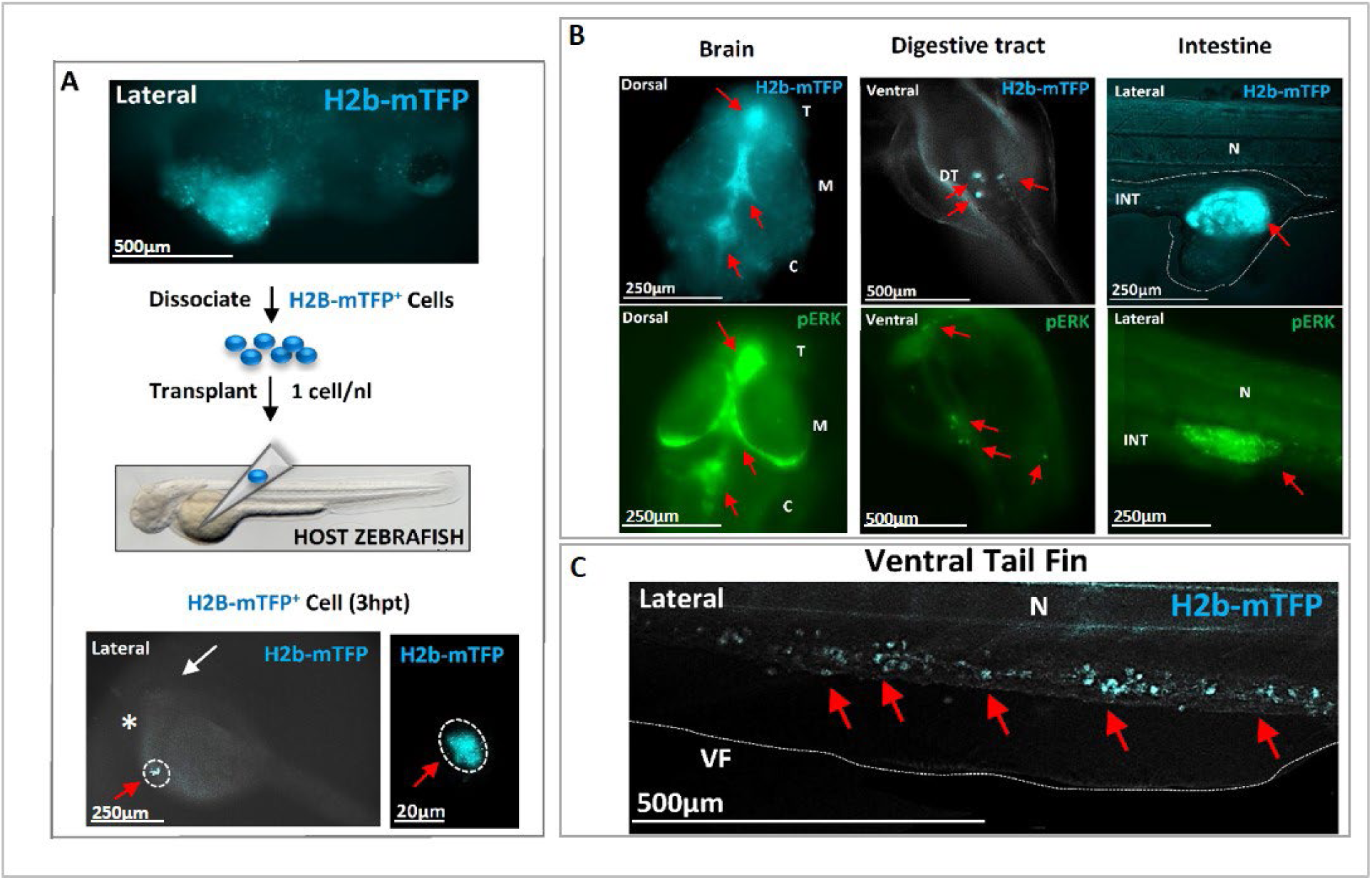
Transplantation of single cell(s) from hyperplasic tissue reveals cancer-initiating potential. **(A)** Following KRASG12V *plus* Ventx activation, hyperplasic tissues (blue fluorescent) are observed at 6dpf, top figure. The cells of the hyperplasia were dissociated and isolated blue (KRAS expressing) cells were transplanted (≈ 1 cell *per* host) at 2dpf in a *Nacre* (*mitf* -/-) zebrafish line for a better tracking of the transplanted cells. The transplanted H2b-mTFP positive (blue) cell (red arrow) can be visualized as early as 3 hours post transplantation (3hpt) in the yolk of the host, close to the duct of Cuvier. White arrow indicates the head/eye (lateral view). **(B)** At 3 dpt the blue cell(s) from the hyperplasic tissue of the donor have colonized the host *Nacre* zebrafish larvae. Tumors in the brain, digestive tract and intestine, are observed and characterized by the blue fluorescence of the donor KRAS expressing cells (red arrows; n=31 out of 52 host individuals). In the bottom, immunofluorescence (IF) analysis of representative host zebrafish larvae with specific high level of phosphorylated ERK activity (pERK, red arrows) in the brain, intestine and digestive tract. **(C)** A high number of exogenous blue fluorescent cells are here observed to migrate in the tail (red arrows). These observations indicate that the transplanted founder cell has both migratory, colonizing behavior, as well as survival growth advantage in the host to form tumors, and thus to re-initiate carcinogenesis. Scale bars and body axes (a: anterior; p: posterior; d; dorsal; v: ventral) are shown; T=telencephalon; M=Mesencephalon; C= Cerebellum; N=notochord; VF= ventral fin.

## DISCUSSION

The surprising frequency of somatic mutations occurring in physiologically normal tissues^40,41^ raises the question of what combinations of events are sufficient for the malignant transformation of a single cell and the rise of the cell of origin of cancer. The robust deterministic process of single-cell cancer induction that we uncovered suggests that the aberrant reactivation of Epi-Driver genes involved in reprogramming/pluripotency (such as VENTX/NANOG, POU5/OCT4) might be relevant to the irreversible malignant transformation of a cell. Reprogramming Epi-Drivers are important regulators of cell viability, survival and proliferation in several cellular contexts, from embryonic stem cells to cancer cells^22–27^. Due to their capacity to modulate epigenetic memory and cell plasticity, these reprogramming factors may drive the early stages of malignant transformation *in vivo* once (re)activated aberrantly. They likely share some mechanism(s) with processes such as induced nuclear-reprogramming^22,23,24^, pluripotency maintenance and/or endogenous cell reprogramming during development^37^. Thus, consistent with our results, the reactivation^37^ of the Neural Crest (NC) Progenitor program (possibly via the stochastic expression of VENTX/NANOG and/or POU5/OCT4) has been shown in BRAF/p53 double mutant cells to yield NC-related tumors *in vivo*^30–32^.We surmise that activation of Epi-Driver genes in NC-related tumors might confer predictable tumorigenesis.

Since cells carrying cancer-causing mutations do not deterministically developed cancers *in vivo*^32,40^, we suggest that the probability of a mutated cell to enter into malignant transformation, or to maintain its physiological functions despite mutations, correlates with the probability of the reactivation of reprogramming factors. Our results support a “Vogelgram”^42^ two-step model (FIG. 5) for irreversible single-cell malignant transformation *in vivo*, mirroring the two hits hypothesis proposed by Berenblum and Shubik^43^.

**FIGURE 5:**
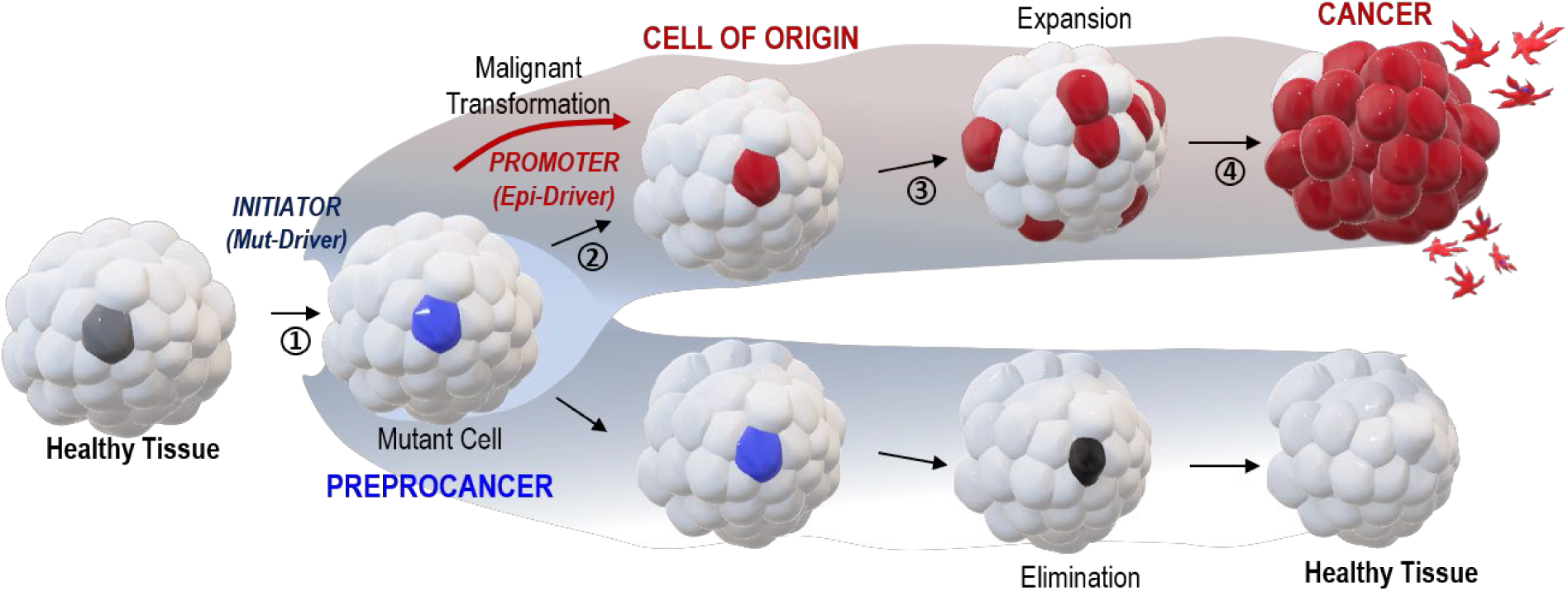
A two-steps “Vogelgram” model of deterministic and irreversible single-cell malignant transformation *in vivo*. Within a healthy tissue, a normal cell (in light grey) acquires (step 1) genetic mutation(s) in driver oncogene(s) (Mut-Drivers such as kRASG12V). Such a mutant cell (in blue, referred as a preprocancer^41^ cell) can (step 2) aberrantly activate the expression of Epi-Drivers involved in pluripotency/reprogramming (e.g. VENTX/NANOG, POU5/OCT4) thus undergoing deterministic and irreversible malignant cell transformation (red cell, the Cell of Origin of Cancer); (step 3) *in situ* short-range dispersal of the early malignant cells and (step 4) further progression to cancer mass and to the appearance of metastatic cells. Inversely, the mutant (preprocancer) cell (in blue) can maintain its physiological functions and be eventually eliminated from the healthy tissue.

Thus, the appearance of Mut-Drivers in a normal (preprocancer^41^) cell^40^ within healthy tissue would act as a permissive but insufficient initiator for malignant cell transformation. Despite the frequency of mutational insults, which are often caused by external cues (chemical carcinogens, UV, etc…) or senescence/ageing, it is known that surveillance mechanisms like cell competition between wild-type and mutated cells safeguard tissue homeostasis and results in active elimination of mutant cells from the tissue^44,45,46^. Our data imply that the aberrant reactivation of Epi-Driver genes involved in reprogramming/pluripotency may promote the irreversible and deterministic malignant transformation in the cell of origin of cancer leading to carcinogenesis.

Our hypothesis is further strengthened by the observation that the variation in cancer risk among tissues can be related to the number of stem cell divisions and tissue renewal^6^. The aberrant reactivation (or maintenance) of Epi-Driver genes in these regenerative events, in cooperation with incident mutations, might be the key deterministic factor in the switching of a normal/healthy cell into the first cell of origin of cancer.

Furthermore, the results reported here are compatible with the recently proposed “ground state theory of cancer initiation7”. According to that theory a malignant transformation may occur in a cell harboring an oncogenic mutation upon a change of its functional state (its “ground state”). Here the transient activation of a reprogramming factor (i.e. Ventx, Nanog or Oct4) possibly synergizes with the oncogene to alter the epigenetic and functional state of the cell allowing its transformation into a tumorigenic cell.

This may also explain the apparent discrepancy between the present deterministic and rapid tumor development upon activation in a single cell of both kRasG12V and a reprogramming factor and the reports of stochastic and random tumor development over weeks following activation of kRasG12V in a whole tissue (brain^47^, pancreas^48^ or intestine^49^). In the latter case, transformed cells may have been triggered to undergo a malignant transformation by the stochastic activation of a reprogramming factor/Epi-Driver, which is consistent with the recent finding that cancer initiation might occurs by transient perturbations of epigenetic regulators^50^.

Importantly, our observation that the progeny of the cell(s) of origin of cancer display a short-range dispersal within the tissue during the earliest phases of tumorigenesis and prior to the effective appearance of tumor mass corroborates a recent theoretical model predicting such an early dispersal during carcinogenesis to explain heterogeneity, therapeutic resistance and tumor relapse^8^.

Being the first experiments to predictably and non-invasively control the malignant transformation of a single cell in an unaltered microenvironment our approach opens a new vista on the study of “the cell of origin of cancer”^51^, an acknowledged enigma of cancer research. As such our results raise many questions: what mechanisms are at work in the transition from the preprocancer state to a tumorigenic state? Is the aberrant reactivation of reprogramming factors a key step in the initiation of carcinogenesis, independent of age, tissue or oncogenic mutation? Do drugs against reprogramming factors reduce the probability of carcinogenesis? Conversely, do some carcinogens (e.g. environmental pollutants, pesticides, endocrine disruptors etc…) act by reactivating those factors?

By allowing for specific and reproductible single cell malignant transformation *in vivo*, our optogenetic approach opens the way for a quantitative study of the initial stages of cancer at the single cell level (e.g. tracking and characterization), that will allow one to address many of these questions.

## MATERIALS&METHODS

### Fish lines and maintenance

Zebrafish were raised and maintained in an approved Fish Facility (C75- 05-32 at IBENS) on a 14–10 h light-dark diurnal cycle with standard culture methods^52^. Embryos collected from natural crosses were staged according to Kimmel^53^. The Tg(*actin*:loxP-EOS-stop-loxP- KRASG12V-T2A-H2B-mTFP) was generated by injecting the plasmid pT24-*actin*:loxP-EOS-stop-loxP- KRASG12V-T2A-H2B-mTFP, which contains the homologous cDNA sequence of KRASG12V from human, with tol2 mRNA transposase. Founder transgenic fish were identified by global expression of Eos. The Tg(*ubi*:Cre-ERT; *myl7*:EGFP) was previously described^54^. The double transgenic line Tg(*actin*:loxP-EOS-stop-loxP-KRASG12V-T2A-H2B-mTFP; *ubi*:Cre-ERT; *myl7*:EGFP) was created by crossing Tg(*actin*:loxP-EOS-stop-loxP-KRASG12V-T2A-H2B-mTFP) and Tg(*ubi*:Cre-ERT; *myl7*:EGFP). Founder double transgenic fish were selected by global expression of Eos and expression of EGFP in the developing heart. Zebrafish were imaged for phenotypic analysis throughout early development, from embryonic to larval stage, and then fixed (at 6dpf) with PAXgene Tissue Container Product (Qiagen) for RT-qPCR or with 4% PFA (Thermofisher) for histological (Hematoxylin & Eosine, H&E) analyses.

### Caged Cyclofen (cCYC) treatment and UV uncaging

To induce kRASG12V expression, 24hpf embryos were incubated in 6μM caged cyclofen (cCYC) for 1 hour (in the dark), rinsed in E3 medium followed by 5 min. photo-activation with a ∼365 nm UV lamp (measured intensity: 4mW/cm^2^). They were then transferred back to E3 medium (with no cCYC) + 10 µM DEX (to release Ventx-GR from its complex with cytoplasmic chaperones). The embryos at 1 dpi were washed 3x in E3 medium to remove DEX. From thereon embryos are exposed neither to cCYC, nor to DEX. For this experiment, we used a benchtop UV lamp (Fisher VL-6-L) which emits a peak wavelength at 365 nm with a FWHM (full width at half maximum) of 40 nm and delivers on the illuminated sample a typical photon flux of ∼1,25 10^-4^ Einstein/(s·m^2^).

Localized uncaging was performed by illumination for 7 min on a Nikon Ti microscope equipped with a light source peaking at 405 nm, Fig.1. The size of the uncaging region was controlled by an iris that defines a circular illumination of diameter ∼ 80 μm. After 0,5-1 h following uncaging the illuminated region of each embryo was imaged to identify and count mTFP positive cell(s) (cells in which the oncogene and its reporter fluorophore was activated). In about 50% of cases one cell was thus photo-activated, the other ∼50% corresponded to more than one activated cell, see Fig.2A. Embryos were transferred to a 12-well plate with E3 + 10 µM DEX, incubated in total darkness overnight and washed 3x with E3 at 1 dpi (to remove DEX).

### Dexamethasone induction

transgenic zebrafish were injected at the one cell stage with the mRNA of a gene-construct Ventx-GR (*Xenopus* ventx2, 450pg), Nanog-GR (mouse Nanog, 100pg) or Pou5/Oct4 (*Xenopus* pou5f3.1/oct91, 100pg) together with a red fluorescent expression marker mRFP (50 pg). The protein products of these constructs are sequestered by cytoplasmic chaperones and released upon incubation of the embryos in a medium containing 10 µM Dexamethasone (DEX). In single cell activation experiments, zebrafish were selected from their strong intensity of fluorescent mRFP signal for local activation of kRasG12V via uncaging of 6µM cCYC (a gift of I.Aujard and L.Jullien).

### Reverse transcriptase quantitative PCR (RT-qPCR)

Zebrafish larvae were fixed (as mentioned above) at 6 dpf. Total RNAs were extracted using the RNeasy micro kit (Qiagen) according to the manufacturer’s protocols. Sample quantity and purity, reverse transcription, pre-amplification and High throughput qPCR were performed as in Zhang et al.^55^. Specifically:

### Reverse transcription

cDNA synthesis was performed using Reverse Transcription Master Mix from Standard Biotools according to the manufacturer’s protocol with random primers in a final volume of 5 μL containing 40 ng total RNA using a Nexus thermocycler (Eppendorf). cDNA samples were diluted by adding 20 μL of low TE buffer [10 mM Tris; 0.1 mM EDTA; pH = 8.0 (TEKNOVA)] and stored at - 20°C.

### Specific target amplification

1.25 μL of each diluted cDNA was used for multiplex pre-amplification with Standard Biotools^®^ PreAmp Master Mix at 18 cycles. In a total volume of 5 μL. the reaction contained 1 μL of pre-amplification mastermix, 1.25 μL of cDNA, 1.25 μL of pooled TaqMan^®^ Gene Expression assays (Life Technologies, ThermoFisher) with a final concentration of each assay of 180 nM (0.2X) and 1 μL of PCR water. The cDNA samples were subjected to pre-amplification following the temperature protocol: 95°C for 2 min, followed by 18 cycles at 95°C for 15 s and 60°C for 4 min. The pre-amplified cDNA were diluted 5X by adding 20 μL of low TE buffer (TEKNOVA) and stored at - 20°C before qPCR.

### High-throughput real time PCR

quantitative PCR was performed using the high-throughput platform BioMark™ HD System and the 48.48 GE Dynamic Arrays (Standard Biotools). 6 μL of sample master mix (SMM) consisted of 1.8 μL of 5X diluted pre-amplified cDNA, 0.3 μL of 20X GE Sample Loading Reagent (Standard Biotools) and 3 μL of TaqMan^®^ Gene Expression PCR Master Mix (Life Technologies, ThermoFisher). Each 6 μL assay master mix (AMM) consisted of 3 μL of TaqMan^®^ Gene Expression assay 20X (Life Technologies) and 3 μL of 2X Assay Loading Reagent (Standard Biotools). 5 μL of each SMM and each AMM premixes were added to the dedicated wells. The samples and assays were mixed inside the chip using MX-FC controller (Standard Biotools). Thermal conditions for qPCR were: 25°C for 30 min and 70°C for 60 min for thermal mix; 50°C for 2 min and 95°C for 10 min for hot start; 40 cycles at 95°C for 15 s and 60°C for 1 min. Data were processed by automatic threshold for each assay, with linear derivative baseline correction using BioMark Real-Time PCR Analysis Software 4.0.1 (Standard Biotools). The quality threshold was set at the default setting of 0.65.

RT-qPCR measurements were done in triplicate on pooled zebrafish larvae and single zebrafish larva (for KR+VX condition), as indicated in the text. Normalization and quantification were obtained with the ΔΔCt method using *rpl13a* as a reference gene. The relative expression of the genes R under the different conditions analysed was calculated as follows using the method described by Livak&Schmittgen^56^: R = 2^-ΔΔCt^ where ΔΔCt = (Ct _gene of interest_ - Ct _reference gene_) _test condition_ - (Ct _gene of interest_ – Ct _reference gene_) _control condition_

### Immunohistochemistry (IHC)

The zebrafish obtained in the various conditions and stages mentioned in the text were fixed in 4% PFA overnight at 4 °C, followed by dehydration with 100% methanol at −20 °C for more than 1 day. After gradual rehydration of methanol and wash with PBS/Tween 0.1%, the embryos were incubated in a blocking solution: 1%Triton, 1% DMSO, 1% BSA and 10% sheep serum (Sigma) in PBS on a shaker for 1 h at room temperature, followed by incubation on a shaker overnight at +4°C with 1:500 anti-phospho-Erk antibody (phospho-p44/42 MAPK (Erk1/2) (Thr202/Tyr204) (D13.14.4E) XP^®^). Rabbit mAb detects endogenous levels of p44 and p42 MAP Kinase (Erk1 and Erk2) when dually phosphorylated at Thr202 and Tyr204 of Erk1 (Thr185 and Tyr187 of Erk2), and singly phosphorylated at Thr202 - this antibody does not cross-react with the corresponding phosphorylated residues of either JNK/SAPK or p38 MAP kinases) (ref 4370S Cell Signalling). After washing with PBS/Tween, embryos were incubated overnight at +4°C with 1/1000 secondary fluorescent conjugated antibody, Donkey anti-Rabbit Alexa Fluor 488 (Thermo Fisher A-21206). To visualize the distribution of Ventx-GR in the fish, embryos were incubated with 1:500 anti-HA antibody, tagging the HA sequence fused to Ventx-GR, as the conjugated antibody (Anti-HA, mouse IgG1, clone 16B12 Alex Fluor^TM^ 488 conjugated, Thermofisher Ref. A21287) on a shaker overnight at 4 °C. All images were taken on a Nikon Ti microscope equipped with a Hamamatsu Orca camera, except for confocal microscopy (Fig.2C, S4B) carried out on a Leica SP5 and Zeiss LSM800 microscope.

### Cell transplantation

Transplantation experiments followed published protocols^57^.Specifically, zebrafish larvae were grown until 6dpi, dissociated in dissociation medium^58^ (025% trypsin-EDTA *plus* 1/10 Collagenase 100mg/ml) 10min at 30°C, mechanically homogenized, resuspended in DMEM-10% Fetal Bovine Serum (FBS), filtered through 70µm nylon mesh (Corning Cell Strainer Ref. 431751) and resuspended in DMEM-10% FBS at 1cell/1nl concentration by using LUNA*-*II cell counter (Logos BioSystem), and injected (∼1 nl) at the level of the yolk sac circulation valley (close to the site of the ducts of Cuvier) of host *nacre* zebrafish at 2 days post-fertilization (2dpf) as shown in Fig.4.

### Microscopy

Fluorescent images were taken on a Nikon Ti microscope equipped with a Hamamatsu ORCA V2+ camera and a 10X plan fluo objective. Filter setting of CFP: excitation at 438 ± 24 nm, emission at 483 ± 32 nm; mEosFP and Alexa 488: excitation at 497 ± 16 nm, emission at 535 ± 22 nm. Image analysis was done using ImageJ software.

## ACKNOWLEDGMENTS

We acknowledge financial support from the ITMO Cancer of Aviesan within the framework of the 2021-2030 Cancer Control Strategy, on funds administered by Inserm. The gene expression analysis was carried out on the high throughput qPCR-HD-Genomic Paris Centre core facility and was supported by grants from the Région Ile de France. We are grateful to I.Aujard and L.Jullien (ENS, Dept. of Chemistry) for a gift of caged cyclofen. We acknowledge useful comments on a draft of this paper by Z.Feng, M.Distel and D.Louvard. P.S. acknowledges useful discussions with C. Rogard, M. Smilla, L.A.M. Scerbo and E.P.A. Scerbo. We thank the Graphic Atelier of Timon Ducos for scientific illustrations (https://timonducos.com/).

## SUPPLEMENTARY FIGURES

**FIGURE S1:**
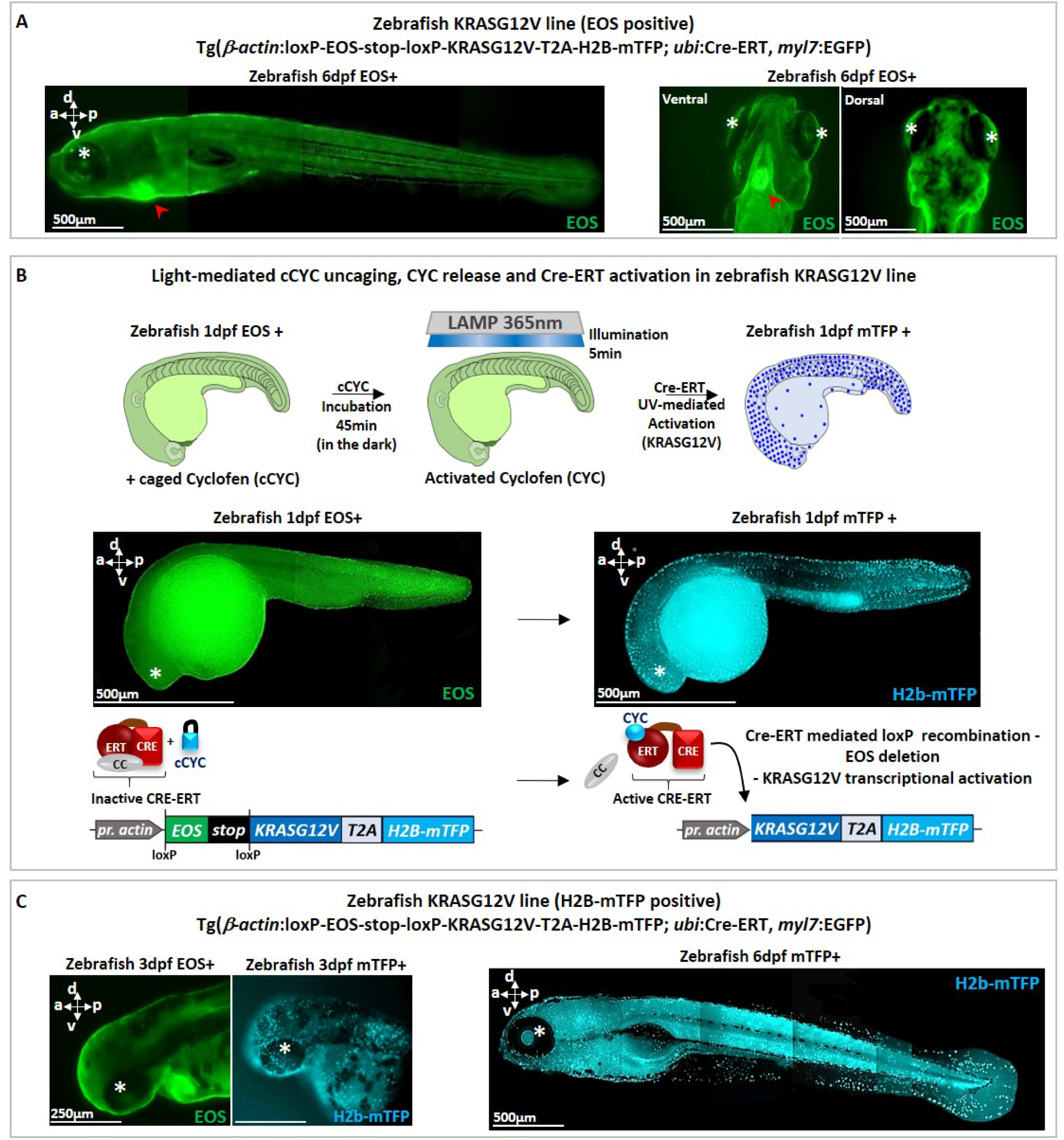
Characterization of the double transgenic line Tg(β−*actin*:loxP-EOS-stop-loxP- KRASG12V-T2A-H2B-mTFP; *ubi*:Cre-ERT; *myl7*:EGFP). **(A)** A transgenic fish at 6dpf displays green EOS fluorescence throughout the body and a strong green fluorescence in the heart due to the additive expression of EGFP under a heart specific (*myl7*) promotor, red arrowhead. **(B)** Upon global illumination (at 1dpf) with a ∼365nm UV lamp in presence of cCYC, cyclofen is uncaged which releases CRE-ERT from its complex with cytoplasmic chaperones and results in floxing of EOS and ubiquitous expression of KRASG12V and H2B-mTFP. The embryos thus display blue fluorescence in the nuclei shown here at 1h post induction (hpi). (**C**) Stable and ubiquitous expression of the blue H2B-mTFP fluorescent protein in zebrafish late embryos (3dpf) and larvae (6dpf). Note that in all pictures the asterisk (*) indicate the eye. Scale bars and body axes (a: anterior; p: posterior; d; dorsal; v: ventral) are shown.

**FIGURE S2:**
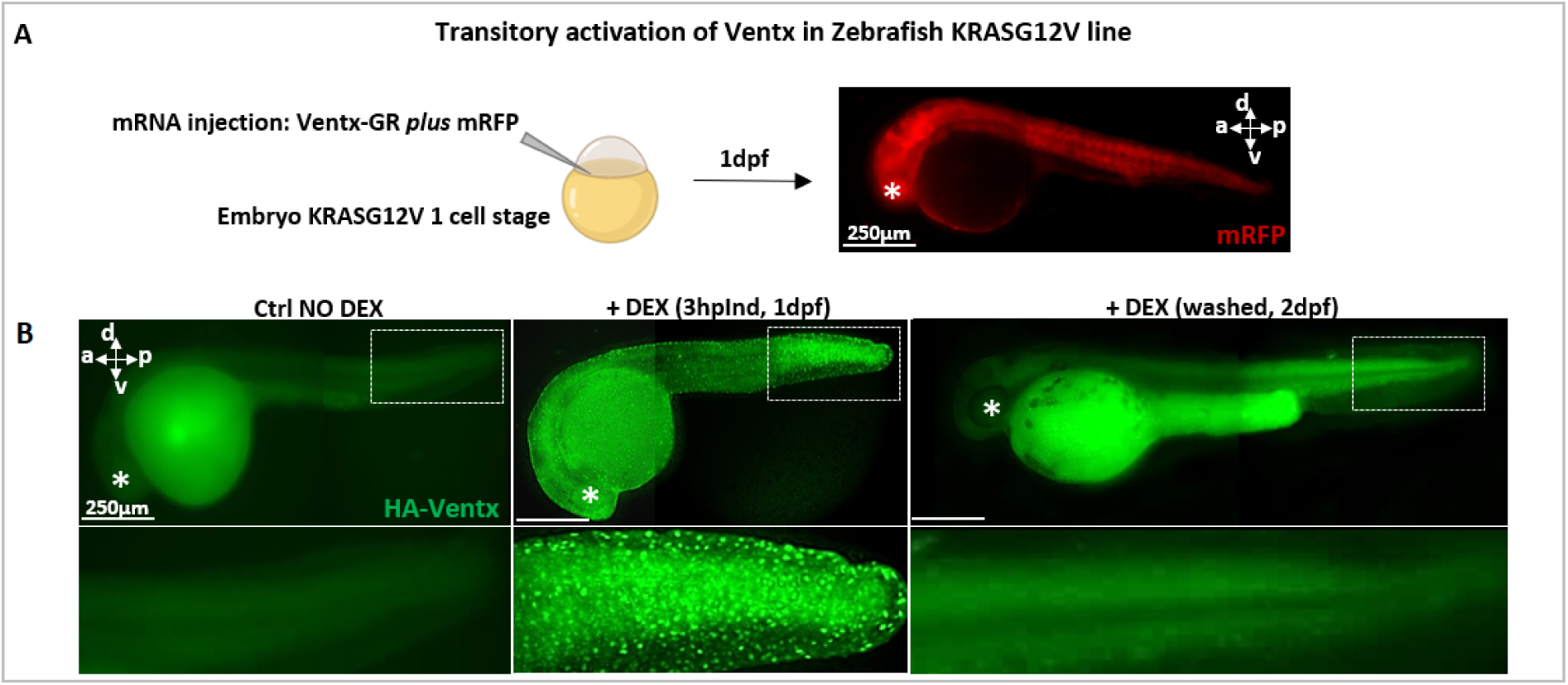
Transient activation of Ventx reprogramming factor. (A) top: At one cell stage Ventx-GR mRNA is injected together with mRFP mRNA (used as a tracer), which expression throughout the embryo (here shown at 1dpf) is a proxy for Ventx-GR expression. **(B) Bottom:** Immunofluorescence analysis show that Ventx protein can be visualized in fixed embryos with the help of an antibody (Ab) directed against the HA tag linking Ventx and GR (and visualized by a second Ab linked to a green fluorescent marker). Upon addition of DEX at 1dpf embryos, Ventx is released from its complex with cytoplasmic chaperones and diffuses into the cell nucleus, resulting in a pointillist image of the nuclei (middle). At 2dpf, after washing out DEX, Ventx protein is no longer detected and the pointillist image is lost (right). Note that in all pictures the asterisk (*) indicate the eye. Scale bars and body axes (a: anterior; p: posterior; d; dorsal; v: ventral; l: left; r: right) are shown.

**FIGURE S3:**
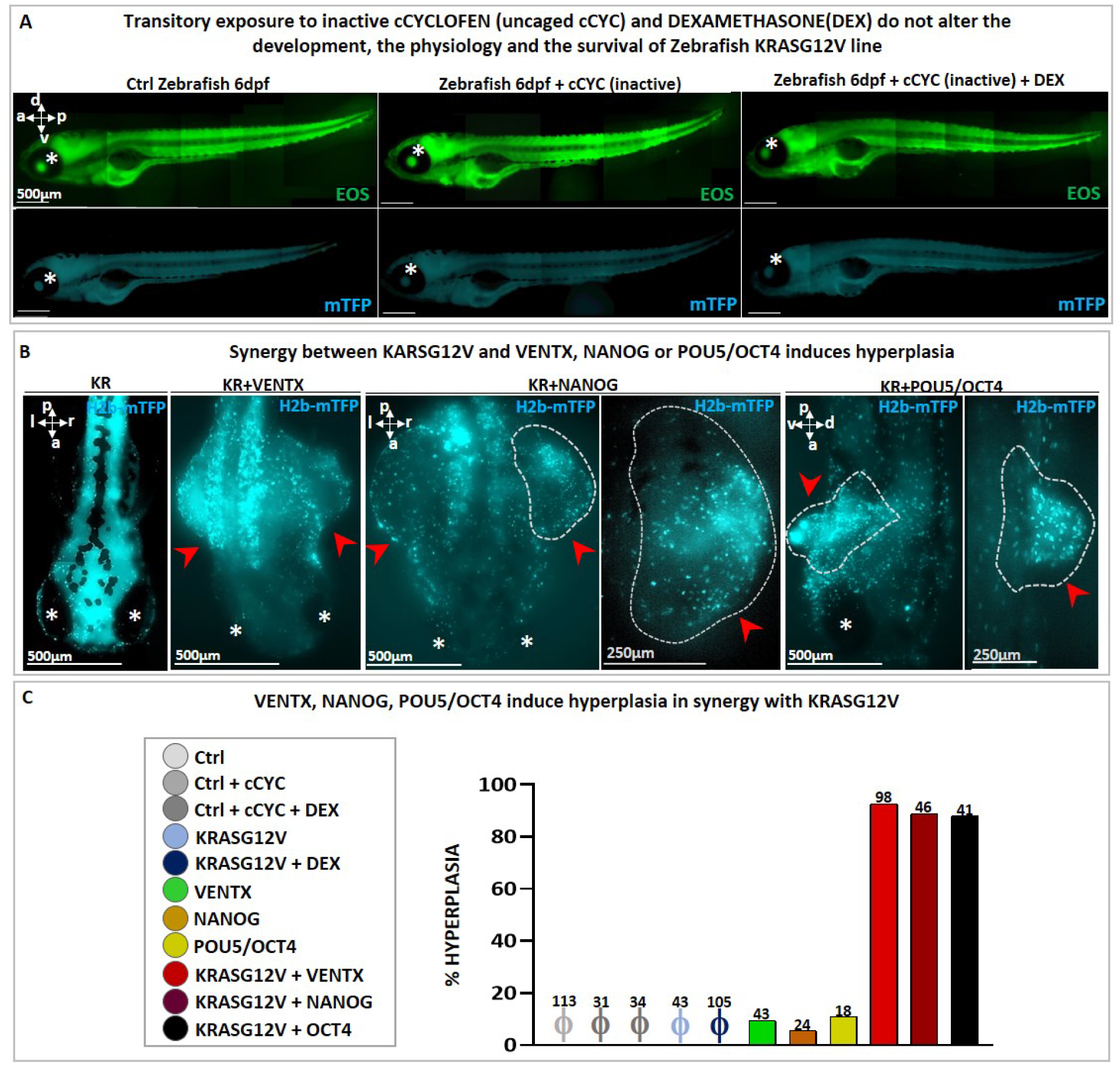
Synergy between a reprogramming factor and KRASG12V oncogene efficiently induces tumors in zebrafish larvae. **(A)** Images of control zebrafish larvae at 6dpf: without any treatment (left), with transient incubation in cCYC (without UV, middle) and with incubation in DEX + cCYC (without UV, right). Note that zebrafish does show neither morphological anomalies nor mortality. **(B)** Global photoactivation (*via* cCYC uncaging with 5 min. UV illumination and Cre-ERT activation) of KRASG12V (blue nuclei) in 1dpf zebrafish without (left) or with subsequent transient activation (right) of Ventx-GR by Dexamethasone (DEX) (described in Fig. S2). These larvae (labelled as KR+VX) develop hyperplasic tissues within 6-9 dpf (dorsal view, red arrows indicate hyperplasia). Global photoactivation (*via* cCYC uncaging and Cre-ERT activation) of KRASG12V (blue nuclei) in 1dpf zebrafish with subsequent transient activation of NANOG-GR or POU5/OCT4- GR by Dexamethasone (DEX) (set-up described in A). These larvae (labelled as KR+NANOG or KR+POU5/OCT4) develop hyperplasic tissues within 6-9 dpf (dorsal view, red arrows indicate hyperplasia). **(C)** Quantification of zebrafish developing abnormal hyperplasia upon the indicated treatments. Note that only the synergy between reprogramming factors (i.e. VENTX, NANOG or POU5/OCT4) and the KRASG12V oncogene reproducibly and efficiently induce tumors in zebrafish larvae. The total number of zebrafish larvae analyzed are indicated above the bars. ϕ indicates no event (0%). Note that in all pictures the asterisk (*) indicate the eye. Scale bars and body axes (a: anterior; p: posterior; d; dorsal; v: ventral; l: left; r: right) are shown.

**FIGURE S4:**
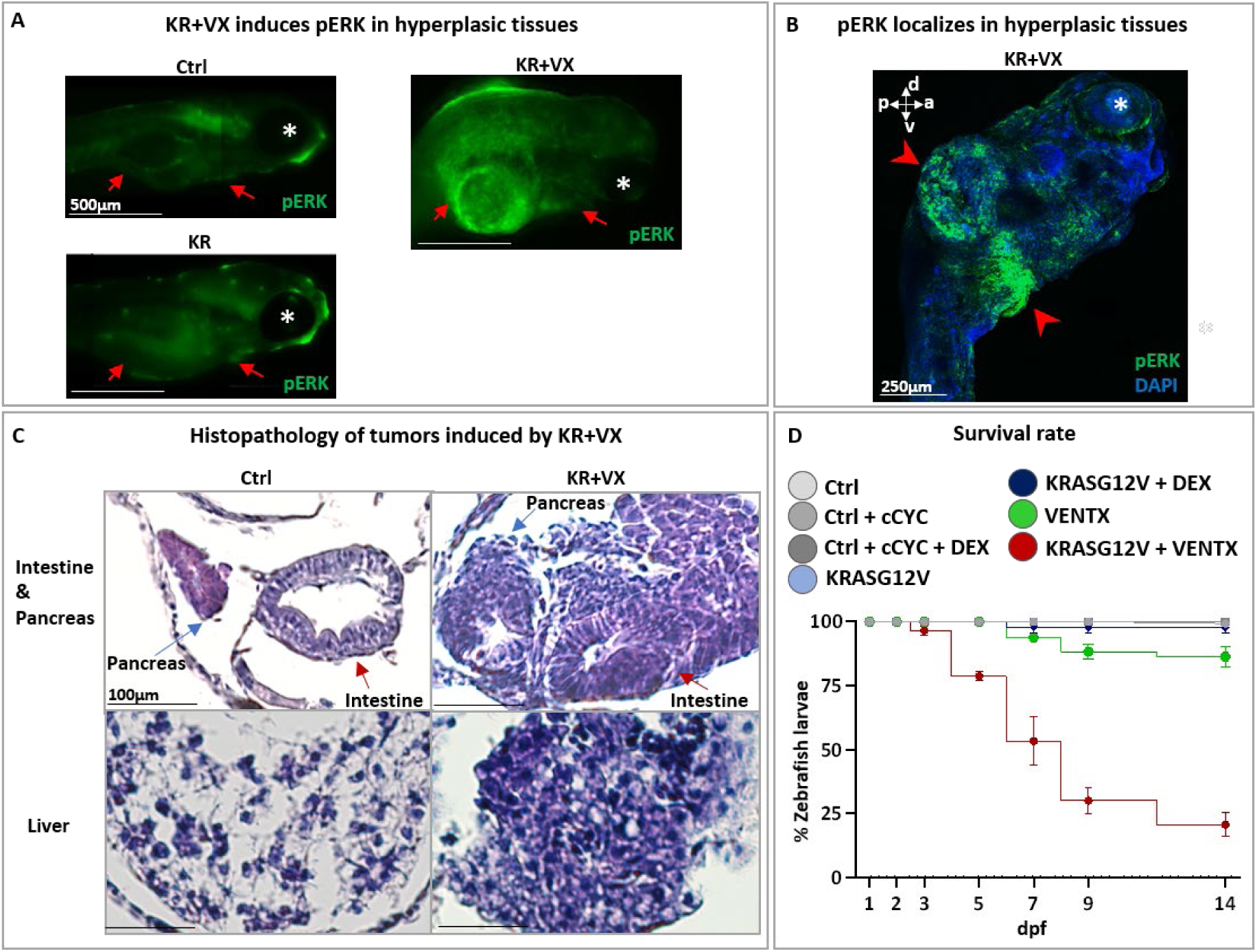
Synergy between the Ventx reprogramming factor and the KRASG12V oncogene induces tumors in zebrafish larvae. **(A)** Phosphorylated ERK (pERK) immunofluorescence at 6dpf in non-activated larvae (Ctrl), in larvea in which only KRASG12V was activated (KR) and in larvae in which both KRASG12V and VENTX were activated globally (KR+VX). Notice in the last one the strong pERK activation in the hyperplasic abdominal outgrowth. The digestive tract is shown between the red arrows. **(B)** Confocal microscopy of KR+VX zebrafish larva at 6dpf (ventro-lateral view) displays the immunofluorescence of active phosphorylated ERK (pERK, in green and indicated by red arrowheads) in the abdomen of a DAPI labelled zebrafish larva. **(C)** Histopathological analysis by Hematoxylin & Eosin (H&E) staining of hyperplasic tissues in zebrafish larvae (6dpf) upon activation of KRASG12V *and* Ventx (KR+VX) is compared with normal tissues (Ctrl). Larvae overexpressing KR+VX develop hyperplasic and dysplastic cancer-like features and tissue outgrowth in the gut/intestine (red arrow), pancreas (blue arrow) and liver. **(D)** Such cancer-like features specifically and exclusively reduce the survival rate to 20% two weeks after induction.

**FIGURE S5:**
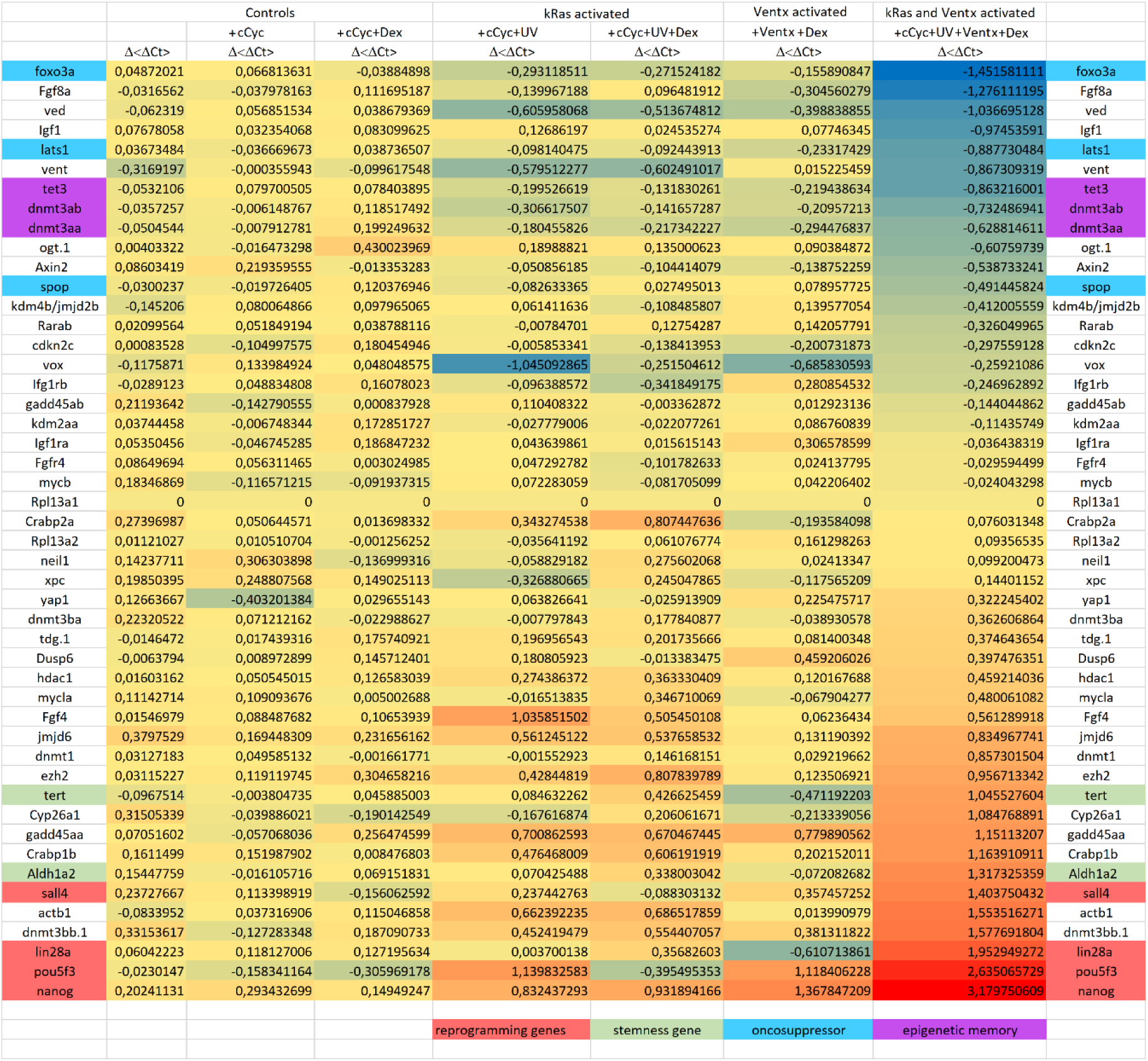
KRASG12V *plus* Ventx induces a cancer-like gene expression signature. Both KRASG12V *and* Ventx-GR (KR+VX) were activated globally in zebrafish at 1dpf. The larvae (n=24) were collected at 6 dpf and processed for RT-qPCR analysis. Compared to the controls (Ctrl; Ctrl+ cCyc; Ctrl + cCYC + Dex) and larvae in which only KRASG12V (+cCyc+UV; +cCyc+UV+ Dex) or Ventx (Ventx+Dex) were activated, the co-activation of KRASG12V + Ventx (+cCyc+UV+VentX+Dex) displayed significant changes in gene expression, with over-expression of genes involved in pluripotency/reprogramming (nanog, pou5f3/oct4, lin28) and stemness (aldh1a, tert) and under-expression of oncosuppressor genes (foxo3, lats1,spop) and genes involved in epigenetic memory (tet3, dnmt).

**FIGURE S6:**
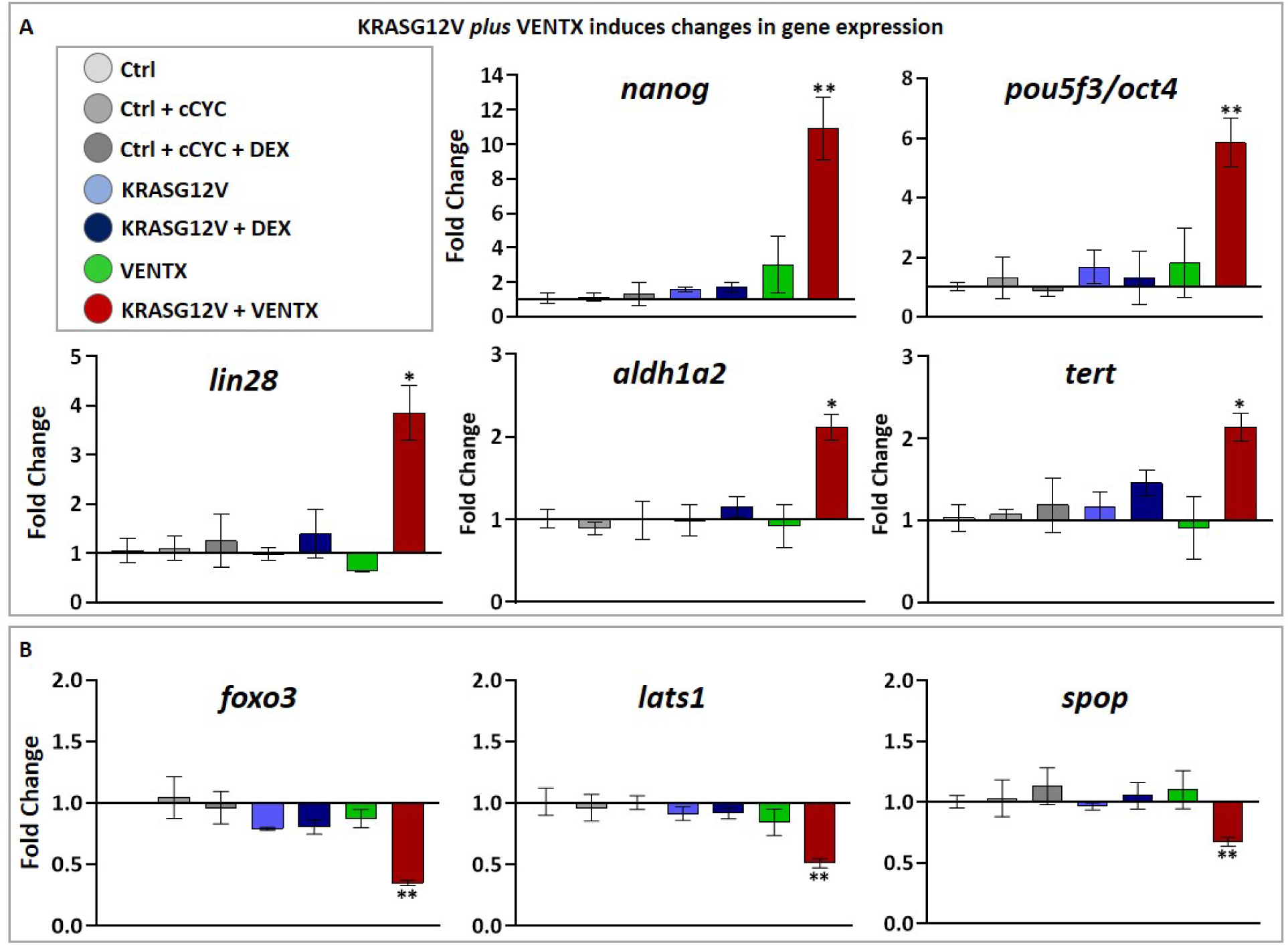
KRASG12V *plus* Ventx induces a cancer-like gene expression signature. Some of the data shown in Fig.S5 are here displayed with their error bars. Compared to the controls (Ctrl in light grey; Ctrl+ cCyc in grey; Ctrl + cCYC + DEX in dark grey) and larvae in which only KRASG12V (KRASG12V in light blue; KRASG12V + DEX in blue) or Ventx (VENTX in green) were activated, the co-activation of KRASG12V + VENTX (in red) displayed significant changes in gene expression. **(A)** Pluripotency/reprogramming factors like *nanog*, *pou5f3*/*oct4* and *lin28,* as well as stemness markers like *aldh1a2* and *telomerase* (*tert*) are significantly up-regulated (p<0.05, compared to Ctrl) by KR+VX but not by KRASG12V (KR) or Ventx (VX) alone. **(B)** Conversely, onco-suppressors such as *foxo3, lats1* and *spop* are significantly down-regulated (p<0.05) by KR+VX but not by KRASG12V (KR) or Ventx (VX) alone when compared to control. For all qPCR graphs, error bars represent s.e.m. values. For statistical analyses, samples were compared with the respective control by Unpaired Student’s t-test. *p<0.05, **p<0.005. ***p<0.0001.

**FIGURE S7:**
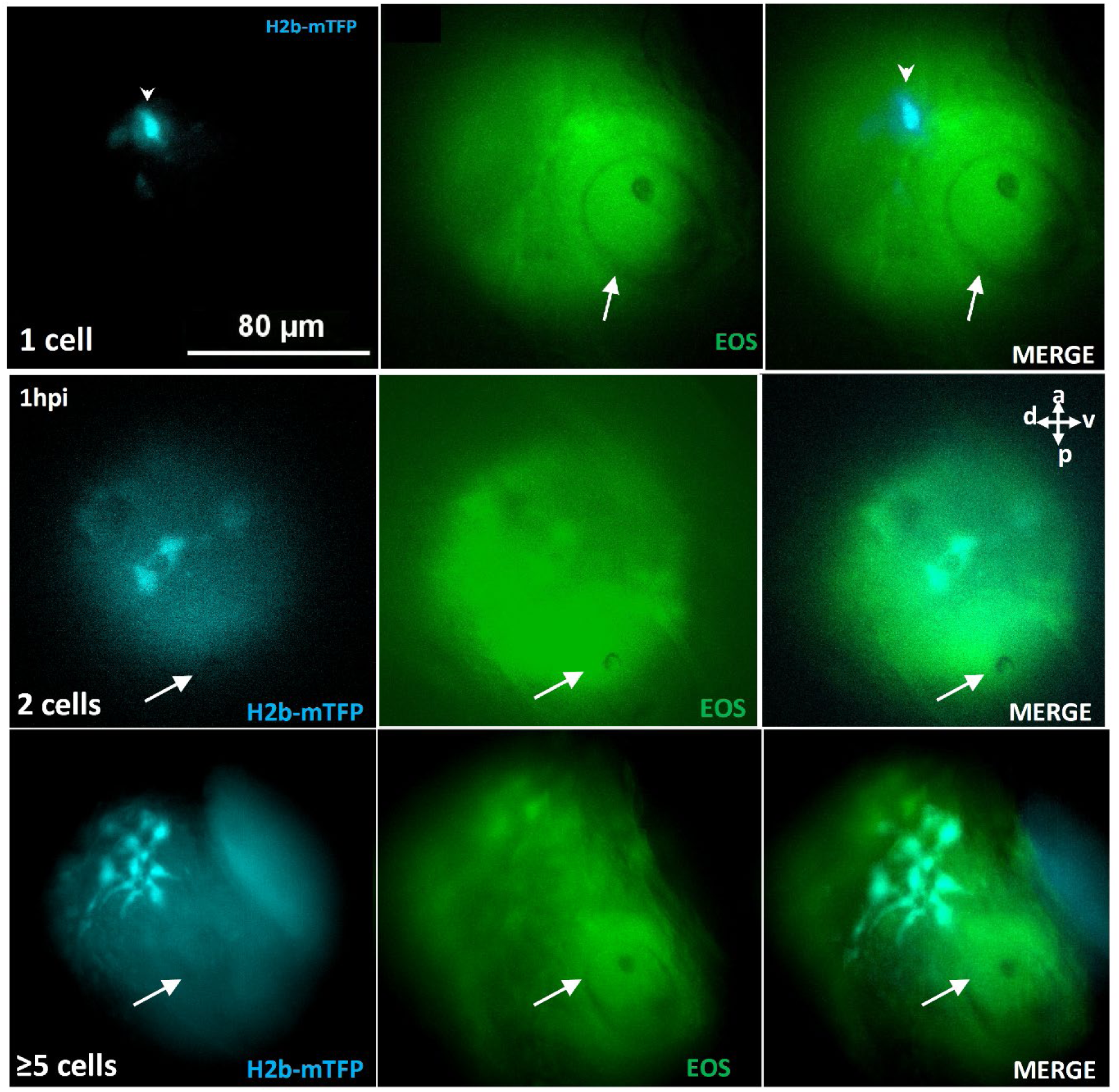
Photo-activation of one or more cells by cCyc uncaging. At 1dpf, an area of diameter ∼80 μm in the vicinity of the otic vesicle (white arrow) was illuminated for 7 min at 405 nm, uncaging cCyc and resulting in one or a few cells in the illuminated region to express the oncogene kRASG12V and its blue fluorescent marker (H2B-mTFP). In these conditions single cell activation represent about 50% of the cases (see Fig.2).

**FIGURE S8:**
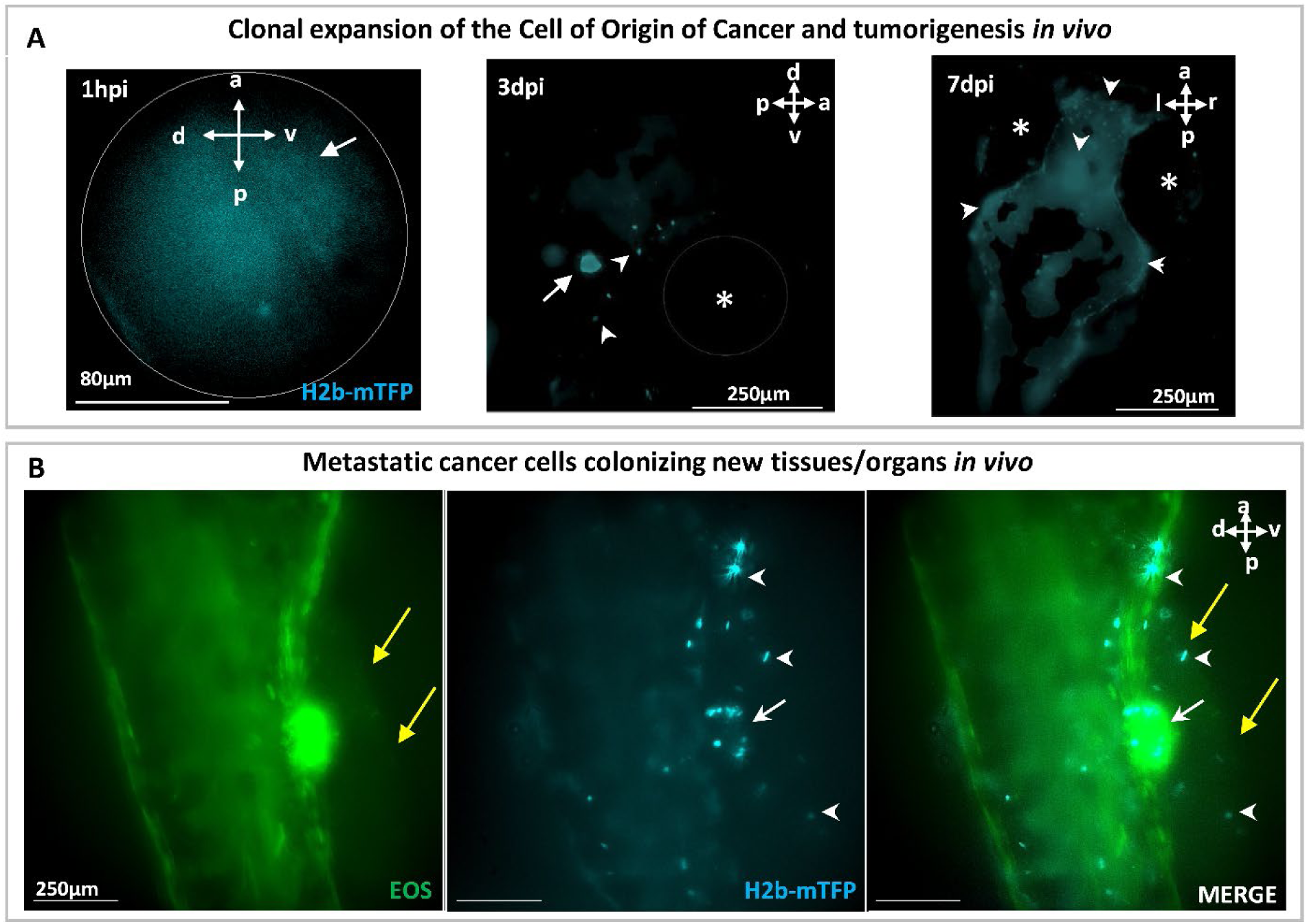
Tracking of the clonal expansion of a photo-induced cell in one embryo over 7 days. **(A)** At 1dpf, a single cell (white arrowhead) in the vicinity of the otic vesicle (white arrow) was photo-induced to express the oncogene KRASG12V, identified within ∼1h (1hpi) by the fluorescent H2B-mTFP marker. Transient (24h) DEX activation of Ventx-GR was done following photoactivation resulting in malignant transformation of the induced cell and clonal expansion. At 3dpi a few cells (white arrowheads) progeny of the induced one are observed in the vicinity of the otic vesicle (white arrow). At 7dpi, a tumor mass is observed in the brain (white arrowheads) and some of the cells (bottom panel) have metastasized (white arrowheads) to colonize new tissues as as the proctodeum, close to the ventral fin (yellow arrows). Body axes are shown (a: anterior;p:posterior; d:dorsal; v:ventral;l:left;r:right).

**FIGURE S9:**
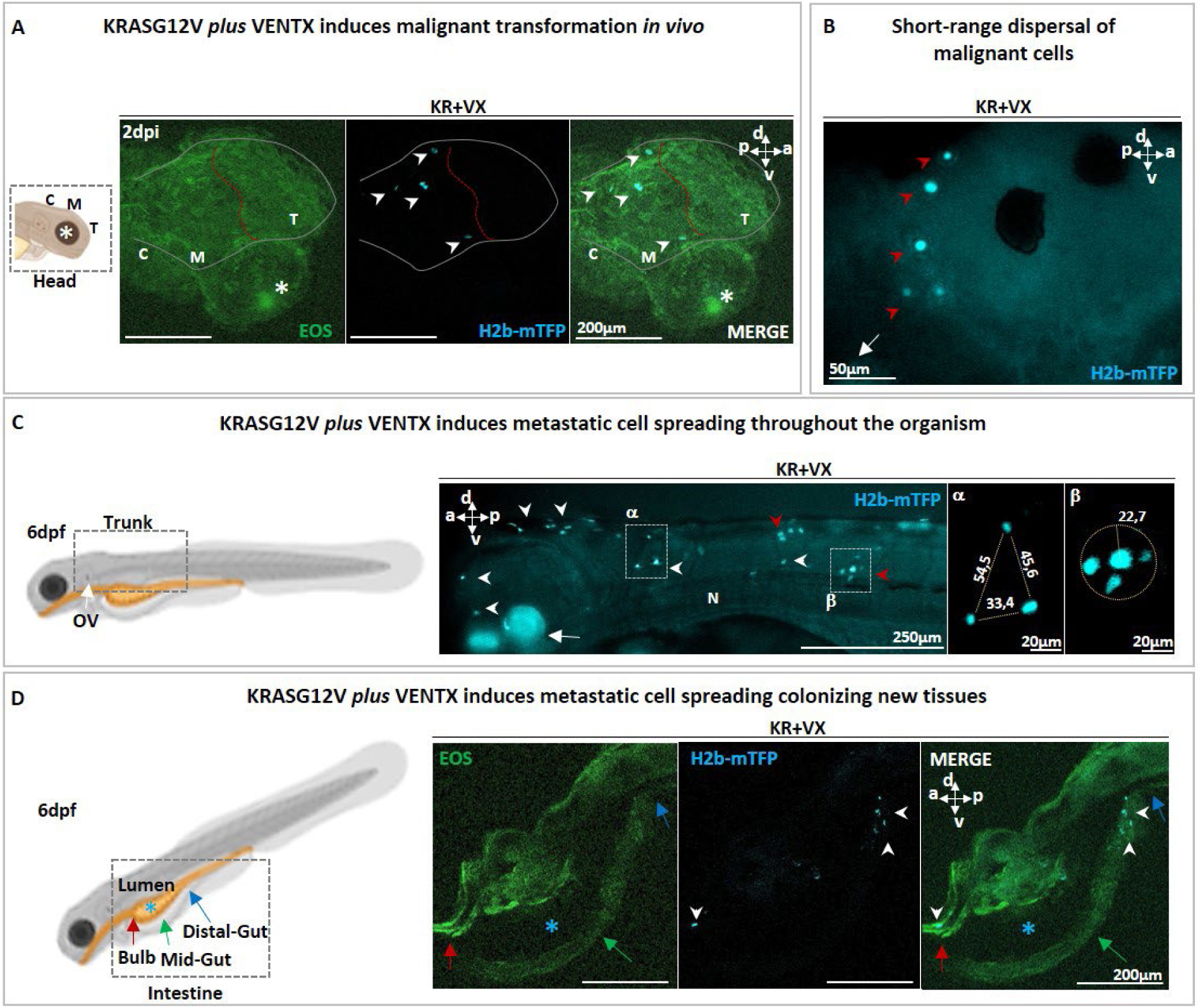
Metastatic spreading following activation of KRASG12V and Ventx (KR+VX). **(A)** Schematic representation and confocal microscopy of zebrafish head showing the early malignant cells (H2B- mTFP ^+^ blue cells; white arrowheads) in the brain. Green EOS fluorescence has been used to visualize the whole zebrafish head. **(B)** Short range dispersal in the brain of the early progeny of the initial malignant cell (H2B-mTFP ^+^ blue cells; red arrowheads). An asterisk (*) indicate the eye and a white arrow the otic vesicle. **(C)** Schematic representation of the zebrafish larva trunk (left, dotted square), where H2B-mTFP^+^ cells (white arrowheads in center panel) can localize in KR+VX larvae, both as isolated cells (as in the dotted square α, 44,5µm of mean distance) or as small clusters (as in the dotted square β, r=22.7µm). **(D)** Schematic representation of zebrafish larva digestive tract (left, dotted square), and confocal microscopy showing H2b-mTFP^+^ cells (white arrowheads, center and right panel) localized in the gut. Green EOS fluorescence has been used to visualize the digestive tract. Red arrows indicate the bulb, the green arrows point to the mid-gut, the blue arrows the distal-gut and the blue asterisk (*) indicate to the gut lumen. Scale bars and body axes (a: anterior; p: posterior; d; dorsal; v: ventral) are shown.

**FIGURE S10:**
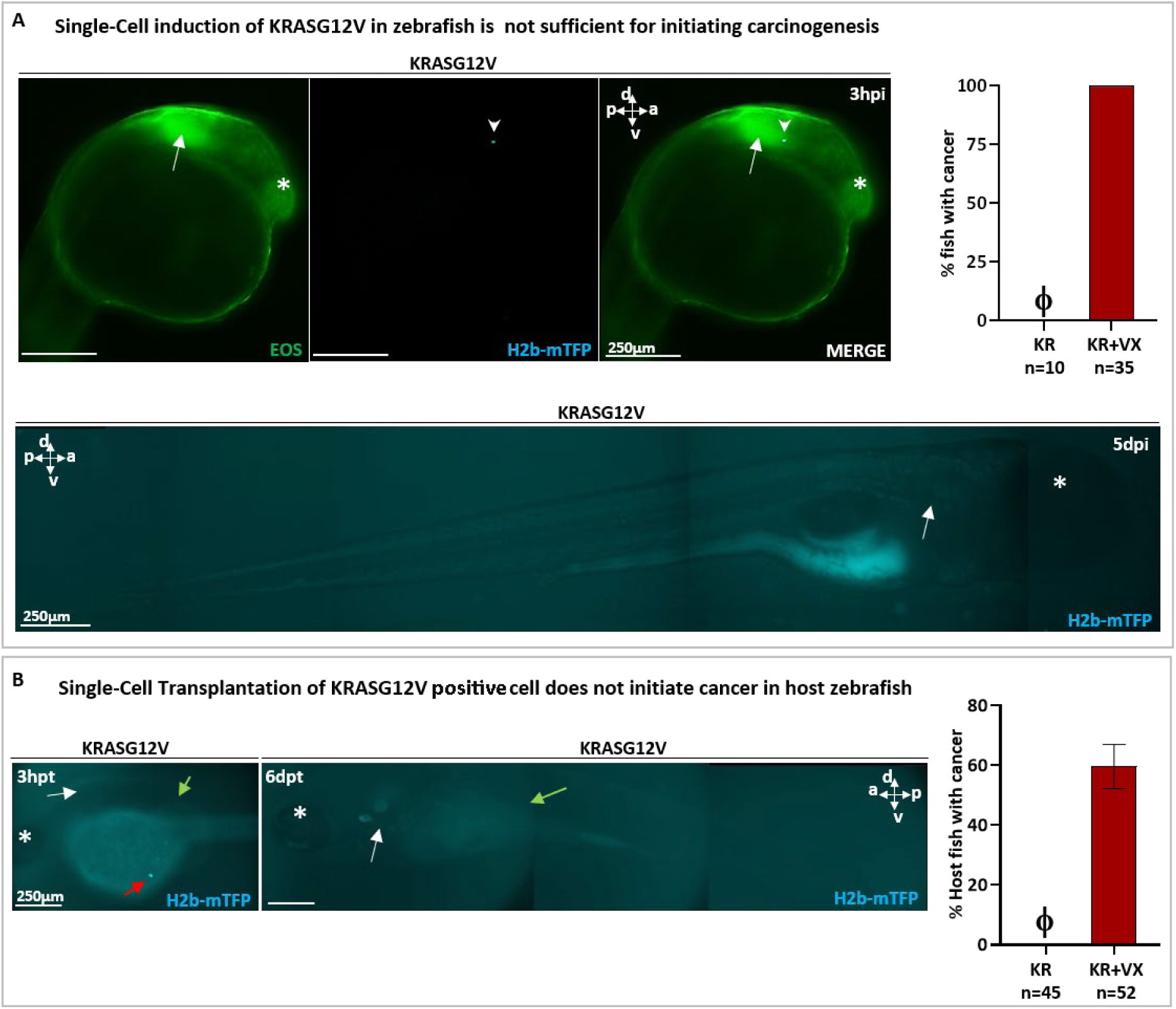
Single-cell activation of the KRASG12V oncogene is not sufficient to initiate carcinogenesis. **(A)** At 1dpf, a single cell in the brain was photo-induced to express the oncogene kRasG12V and identified (white arrowhead) within ∼30 min by the blue fluorescence of the expression marker (H2b-mTFP). The otic vesicle (indicated by white arrow) is used as a spatial reference. At 5 days post induction (5 dpi), the activated cell has disappeared. Note in the graph that, whereas KRASG12V *plus* VENTX (KR+VX) efficiently induces cancer (100%), KRASG12V alone (KR) is not sufficient (0%, indicated by ϕ). **(B)** Expression of kRasG12V was activated in 1 dpf embryos (without subsequent activation of Ventx). The cells of the larvae were dissociated and isolated cells (kRas expressing, H2b-mTFP^+^ blue cells) were transplanted (≈ 1 cell *per* host) at 2dpf in a *Nacre* (*mitf* -/-) zebrafish line for a better tracking of the transplanted cells. The transplanted H2b-mTFP^+^ blue cell (red arrow) can be visualized as early as 3 hours post transplantation (3hpt) in the yolk of the host. At 6 dpt the blue cell has disappeared in host *Nacre* zebrafish larvae. Note in the graph that, whereas the transplanted cell experiencing KRASG12V *plus* VENTX (KR+VX) activation efficiently give rise to cancer (60%), KRASG12V alone (KR) is not sufficient to initiate cancer in host zebrafish (0%, indicated by ϕ). In all figures, the otic vesicle (white arrow) is indicated as well as the eye (white asterisk: *). Scale bars and body axes (a: anterior; p: posterior; d; dorsal; v: ventral; l: left; r: right) are shown.

## Notes

### Competing Interest Statement

The authors have declared no competing interest.

### Summary of Updates

- New Figures - New Supplementary Data

## REFERENCES

1 Cairns, J. Mutation selection and the natural history of cancer, Nature 255, 197–200 (1975)

2 Riva, L., et al. The mutational signature profile of known and suspected human carcinogens in mice. Nat. Genet. 52, 1189–1197 (2020).

3 Adashek JJ, Kato S, Lippman SM, Kurzrock R. The paradox of cancer genes in non-malignant conditions: implications for precision medicine. Genome Med. 12, 16 (2020)

4 Hanahan, D. & Weinberg, R. A., The hallmarks of cancer, Cell 100, 57–70 (2000).

5 Soto, A. M. & Sonnenschein, C. The tissue organization field theory of cancer: a testable replacement for the somatic mutation theory. BioEssays 33, 332–340 (2011).

6 Tomasetti C, Vogelstein B, Cancer etiology. Variation in cancer risk among tissues can be explained by the number of stem cell divisions. Science 347, 78–81 (2015).

7 Jassim A, Rahrmann E.P, Simons B.D. and Gilbertson R.J., Cancers make their own luck: theories of cancer origins, Nat.Rev.Can. 23, 710–724 (2023)

8 Waclaw B. et al., A spatial model predicts that dispersal and cell turnover limit intratumor heterogeneity, Nature 25, 261–264 (2015)

9 Cairns, J., Mutation selection and the natural history of cancer, Nature 255, 197–200 (1975)

10 Nowell, P. C., The clonal evolution of tumor cell populations, Science 194, 23–28 (1976).

11 Hanahan, D. & Weinberg, R. A., The hallmarks of cancer, Cell 100, 57–70 (2000).

12 Tomasetti C, Vogelstein B, Cancer etiology. Variation in cancer risk among tissues can be explained by the number of stem cell divisions. Science 347, 78–81 (2015).

13 Frumkin, D et al., Cell lineage analysis of a mouse tumor, Cancer research 68, 5924–5931 (2008).

14 Malumbres M, Barbacid M., RAS oncogenes: the first 30 years, Nat Rev Cancer 3, 459–465 (2003).

15 Pylayeva-Gupta Y, Grabocka E, Bar-Sagi D, RAS oncogenes: weaving a tumorigenic web, Nat Rev Cancer 11, 761–74 (2011).

16 Strachan, T. & Read, A. P., “Chapter 18: Cancer Genetics”. Human molecular genetics 2 (1999, John Wiley, New York).

17 Blasco MA, Telomeres and human disease: ageing, cancer and beyond, Nat.Rev.Genet. 6, 611–622 (2005)

18 Catalogue of Somatic Mutations in Cancer; https://cancer.sanger.ac.uk/cosmic

19 Punekar SR, Velcheti V, Neel BG, Wong KK. The current state of the art and future trends in RAS-targeted cancer therapies. Nat.Rev.Clin.Oncol. 19, 637–655 (2022).

20 Merz V et al., Targeting KRAS: The Elephant in the Room of Epithelial Cancers. Front.Oncol. 11, 638360 (2021).

21 Vogelstein B, Papadopoulos N, Velculescu VE, Zhou S, Diaz LA Jr, Kinzler KW, Cancer genome landscapes. Science, 339, 1546–58 (2013).

22 Hepburn AC et al., The induction of core pluripotency master regulators in cancers defines poor clinical outcomes and treatment resistance. Oncogene 38, 4412–4424 (2019).

23 Villodre ES, Kipper FC, Pereira MB, Lenz G. Roles of OCT4 in tumorigenesis, cancer therapy resistance and prognosis. Cancer Treat. Rev. 51, 1–9 (2016).

24 Ducos B, Bensimon D, Scerbo P. Vertebrate Cell Differentiation, Evolution, and Diseases: The Vertebrate-Specific Developmental Potential Guardians *VENTX*/*NANOG* and *POU5*/*OCT4* Enter the Stage. Cells 11, 2299 (2022).

25 Rawat VP et al., The vent-like homeobox gene VENTX promotes human myeloid differentiation and is highly expressed in acute myeloid leukemia. Proc Natl Acad Sci U S A. 107, 16946–51 (2010).

26 Laise P et al., Developmental and MAPK-responsive transcription factors drive distinct malignant subtypes and genetic dependencies in pancreatic cancer. bioRxiv 2020.10.27.357269;.

27 Park Y et al., Transcriptomic Landscape of Lower Grade Glioma Based on Age-Related Non-Silent Somatic Mutations. Curr. Oncol. 28, 2281–2295 (2021).

28 Shibata H. et al., In vivo reprogramming drives Kras-induced cancer development, Nat.Comm. 9, 2081 (2018).

29 Adashek JJ, Kato S, Lippman SM, Kurzrock R. The paradox of cancer genes in non-malignant conditions: implications for precision medicine. Genome Med. 12, 16 (2020)

30 Baggiolini A et al., Developmental chromatin programs determine oncogenic competence in melanoma. Science 373, eabc1048 (2021).

31 Ablain J et al., Human tumor genomics and zebrafish modelling identify *SPRED1* loss as a driver of mucosal melanoma. Science 362, 1055–1060 (2018).

32 Kaufman CK et al., A zebrafish melanoma model reveals emergence of neural crest identity during melanoma initiation. Science 351, aad2197 (2016).

33 Feng Z, et al., Optical Control of Tumor Induction in the Zebrafish. Sci.Rep. 7, 9195 (2017).

34 Sinha DK et al., Photoactivation of the CreERT2 recombinase for conditional site-specific recombina-tion with high spatio-temporal resolution. Zebrafish 7, 199–204 (2010)

35 Zhang W et al., Control of protein activity and gene expression by cyclofen-OH uncaging, ChemBioChem 19,1–8 (2018).

36 Feil, R., et al. Ligand-activated site-specific recombination in mice. Proc Natl Acad Sci U S A 93, 10887–10890 (1996).

37 Scerbo P, Monsoro-Burq AH. The vertebrate-specific VENTX/NANOG gene empowers neural crest with ectomesenchyme potential. Sci Adv. 6, 1469 (2020)

38 Lam KHB et al., Topographic mapping of the glioblastoma proteome reveals a triple-axis model of intra-tumoral heterogeneity, Nat Commun, 13, 116 (2022).

39 Campbell NR, Cooperation between melanoma cell states promotes metastasis through heterotypic cluster formation. Dev Cell. 56, 2808–2825 (2021)

40 Martincorena I and Campbell PJ., Somatic mutation in cancer and normal cells, Science 349,1483–9 (2015)

41 Brash, D.E., Preprocancer: normal skin harbor cancer causing mutations, Science 349,867–868 (2015)

42 Fearon E. R. and Vogelstein B., A genetic model for colorectal tumorigenesis, Cell 61, 759–767 (1990).

43 Berenblum I. and Shubik P., The persistence of latent tumor cells induced in the mouse’s skin by a single application of 9, 10-dimethyl-1, 2- benzanthracene. Br. J. Cancer 3, 384–386 (1949).

44 Brown S, et al., Correction of aberrant growth preserves tissue homeostasis, Nature 548, 334–337 (2017)

45 Porazinski S, de Navascués J, Yako Y, Hill W, Jones MR, Maddison R, Fujita Y, Hogan C., EphA2 Drives the Segregation of Ras-Transformed Epithelial Cells from Normal Neighbors, Curr Biol. 26, 3220–3229 (2016)

46 van Neerven SM and Vermeulen L., Cell competition in development, homeostasis and cancer, Nat Rev Mol Cell Biol. 24, 221–236 (2023)

47 Ju B., et al., Oncogenic KRAS promotes malignant brain tumors in zebrafish, Mol.Cancer 14, 18 (2015)

48 Park JD and Leach SD, Zebrafish model of KRAS-initiated pancreatic cancer, Ann.Cells&Sys. 22, 353–359 (2018)

49 Lu J-W., et al., Inducible Intestine-Specific Expression of kRasG12V Triggers Intestinal Tumorigenesis In Transgenic Zebrafish, Neoplasia 20, 1187–1197 (2018)

50 Parreno, V., et al., Transient loss of Polycomb components induces an epigenetic cancer fate, Nature, 629,688 (2024).

51 Visvader JE. Cells of origin in cancer. Nature 469, 314–22 (2021)

52 Westerfield, M. The Zebrafish Book. A Guide for The Laboratory Use of Zebrafish (Danio rerio) (2000).

53 Kimmel, C.B., Ballard, W.W., Kimmel, S.R., Ullmann, B., and Schilling, T.F. Stages of embryonic development of the zebrafish. Dev. Dyn. 203, 253–310 (1995)

54 Mosiman, C. et al., Ubiquitous transgene expression and Cre-based recombination driven by the *ubiquitin* promoter in zebrafish, Development 138, 169–177 (2011)

55 W.Zhang, et al., Fgf8 dynamics and critical slowing down may account for the temperature independence of somitogenesis, Commun.Biol. 5, 113 (2022).

56 Livak KJ, Schmittgen TD. Analysis of relative gene expression data using real-time quantitative PCR and the 2^-ΔΔC(T)^ Method, Methods 25, 402–8 (2001)

57 S. Nicoli and M. Presta, The zebrafish/tumor xenograft angiogenesis assay, Nat.Prot. 2, 2918–23 (2007)

58 Bresciani E, E. Broadbridge and P.P. Liu, An efficient dissociation protocol for generation of single cell suspension from zebrafish embryos and larvae, MethodsX 5, 1287–1290 (2018).

